# Robust Identification of Drug Mechanism of Action Associated Genes Across Heterogeneous Drug Treated Datasets

**DOI:** 10.64898/2026.10.15.759469

**Authors:** Mahfuza Sharmin, Maliha Aziz, Ewy Mathe

## Abstract

Identifying genes associated with the mechanism of action (MOA) of anticancer therapies is essential for understanding drug response. Functional genomics approaches, including genome-wide CRISPR knock out screens, enable systematic gene prioritization following treatment; however, findings from individual datasets are typically interpreted after identifying overlapping genes across datasets to confirm biological hypotheses. Integrating such heterogeneous datasets remains challenging due to variability across experimental conditions. Existing approaches can miss biologically relevant genes due to dataset-specific noise, can fail to capture genes exhibiting consistent but moderate effects across studies. Here, we present ConcuRx, a statistical framework for cross-study integration and gene prioritization across heterogeneous CRISPR drug-response screens. This approach produces more stable gene rankings than single-dataset analyses, improving prioritization of known MOA-associated genes and recovering biologically relevant genes that are overlooked by average- and overlap-based approaches. Applied to CRISPR drug-response screens for the synergistic drug pair of ATR and PARP inhibitors, ConcuRx strengthened support for 15 experimentally validated hypotheses collected from published studies, including mechanisms supported by independent studies not included in the ConcuRx analysis. These findings demonstrate that modelling between-study variability improves identification and prioritization of MOA-associated genes and provides a robust strategy for integrating heterogeneous functional genomics data in precision oncology.

**GRAPHICAL ABSTRACT:** 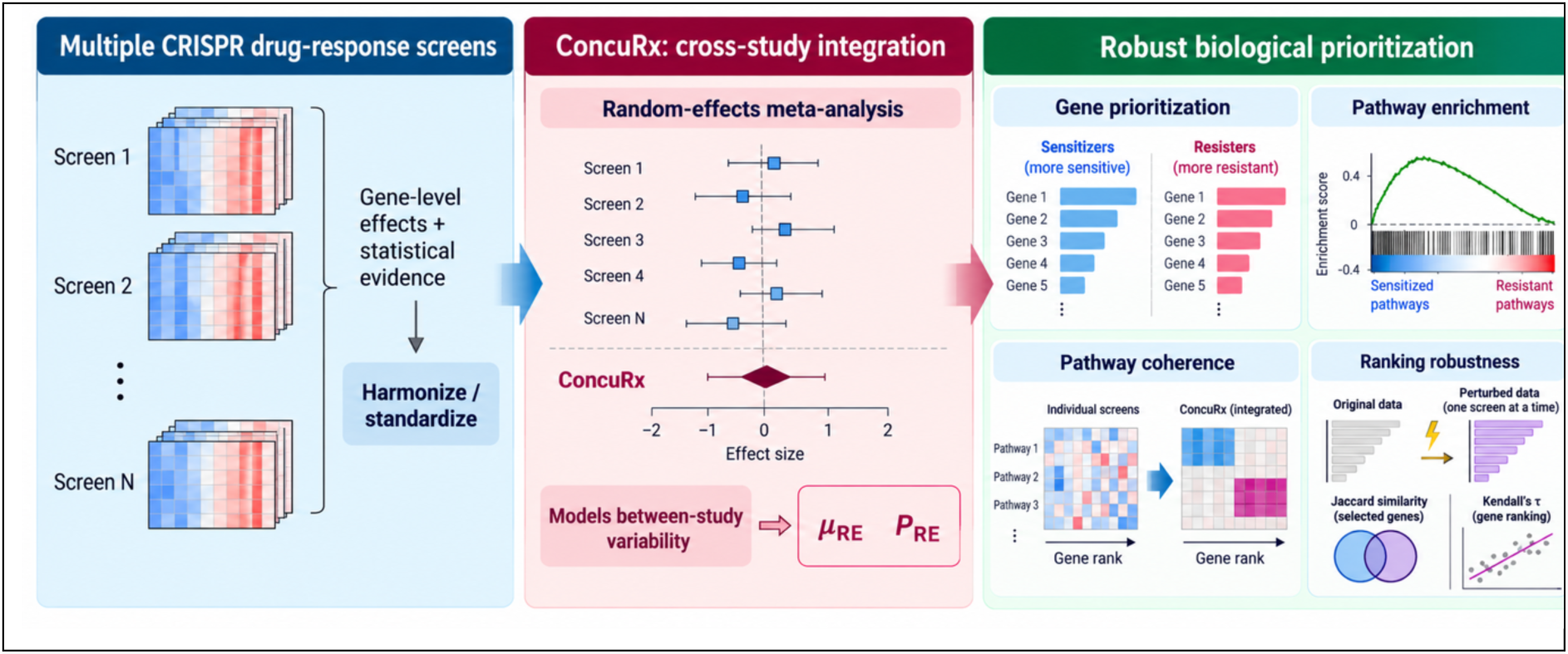

## INTRODUCTION

Identifying genes associated with the mechanism of action (MOA) of anticancer therapies is fundamental to understanding drug response, resistance, and the rational design of combination treatments. Functional genomics approaches, including CRISPR-based genetic screens, have enabled systematic interrogation of gene function and pathway activity in cells with specific disease. These technologies provide complementary views of cellular response to perturbation and are increasingly used to prioritize genes linked to drug mechanisms.

In practice, findings from such studies are often evaluated across independent datasets (e.g. across drugs of same class, across cell lines) to assess reproducibility. A common strategy is to identify genes that overlap between studies through intersection-based analyses or to aggregate effect sizes by averaging across datasets^1–3^. While straightforward, overlap-based approaches are inherently conservative and can discard substantial biological signal. Genes with strong but nonsignificant effects in individual datasets, genes affected by dataset-specific noise, or genes exhibiting consistent but moderate effects across studies may be excluded, despite their potential relevance to underlying biology. Simple averaging retains more information but does not incorporate uncertainty associated with individual estimates or explicitly account for between-study variability, often requiring arbitrary thresholds to prioritize genes.

These limitations are particularly pronounced in the context of heterogeneous functional genomics data. In transcriptomic studies, integration across datasets is often addressed through batch correction and normalization frameworks that aim to align measurements across studies^4,5^. Such approaches are less directly applicable to drug-treated CRISPR knockout screens, in which gene-perturbation effects depend on experimental design, guide-library composition, treatment conditions, and cellular context. Although normalization and bias correction are integral to the analysis of individual CRISPR screens, differences among independent screens complicate direct cross-study integration.

Beyond these technical challenges, datasets are often generated across distinct biological contexts, including different cell lines, genetic or mutational backgrounds, drug concentrations, and related but nonidentical drug perturbations. These differences can represent meaningful biological variation rather than unwanted technical noise; therefore, treating them as batch effects and directly normalizing or aligning datasets may remove or obscure context-specific drug–gene dependencies. In contrast, random-effects meta-analysis^6^ does not require removal of these between-study differences but explicitly models their variability while estimating an overall gene effect. As a result, cross-study integration of CRISPR knockout screens frequently relies on analyzing datasets independently or comparing results through overlap-based strategies^7–10^, without explicitly modelling between-study variability.

Here, we present ConcuRx, a statistical framework for cross-study integration and gene prioritization across heterogeneous CRISPR drug-response screens. ConcuRx formulates cross-study gene prioritization as a random-effects meta-analysis problem and uses the Paule–Mandel estimator ^11–13^ to accommodate the limited number of screens commonly available for individual drugs. Rather than requiring agreement among top-ranked genes, ConcuRx integrates gene-level effect magnitude and uncertainty while explicitly modeling between-study heterogeneity. This enables aggregation of signals that may be weak, variable, or noisy in individual screens while retaining information about cross-study variability. Although random-effects meta-analysis is well established ^14^, its use as a framework for cross-study gene prioritization provides a systematic alternative to overlap- and average-based integration of heterogeneous CRISPR drug-response screens ^15–19^.

We demonstrate ConcuRx using CRISPR screens for ATR and PARP inhibitors, a synergistic drug pair with well-characterized genetic dependencies and mechanisms of response. Across these datasets, ConcuRx recovers biologically relevant genes missed by conventional approaches, captures coherent pathway-level signals, improves prioritization of MOA-associated genes in the presence of cross-study variability, and produces more stable gene rankings under data perturbation. We further evaluate ConcuRx against 15 experimentally supported hypotheses from 15 published studies, including mechanisms derived from independent studies whose datasets were not included in the ConcuRx analysis. Of these, eight hypotheses are discussed in the main text, while the remaining seven (H5, H8, and H11–H15) are presented in the Supplementary Materials (Figure S8-11). Together, these analyses establish ConcuRx as a robust framework for integrating heterogeneous functional-genomics screens and prioritizing reproducible MOA-associated genetic dependencies while retaining biologically meaningful cross-study variability.

## MATERIAL AND METHODS

### ConcuRx framework

We developed ConcuRx, a computational framework for integrating gene-level effects across heterogeneous functional-genomics screens. ConcuRx harmonizes gene-level results from individual screens and integrates them using random-effects meta-analysis to generate cross-study effect estimates and gene rankings (Figure 1A). Data processing, statistical integration, and downstream validation were implemented using Python and R as described below.

**Figure 1.**
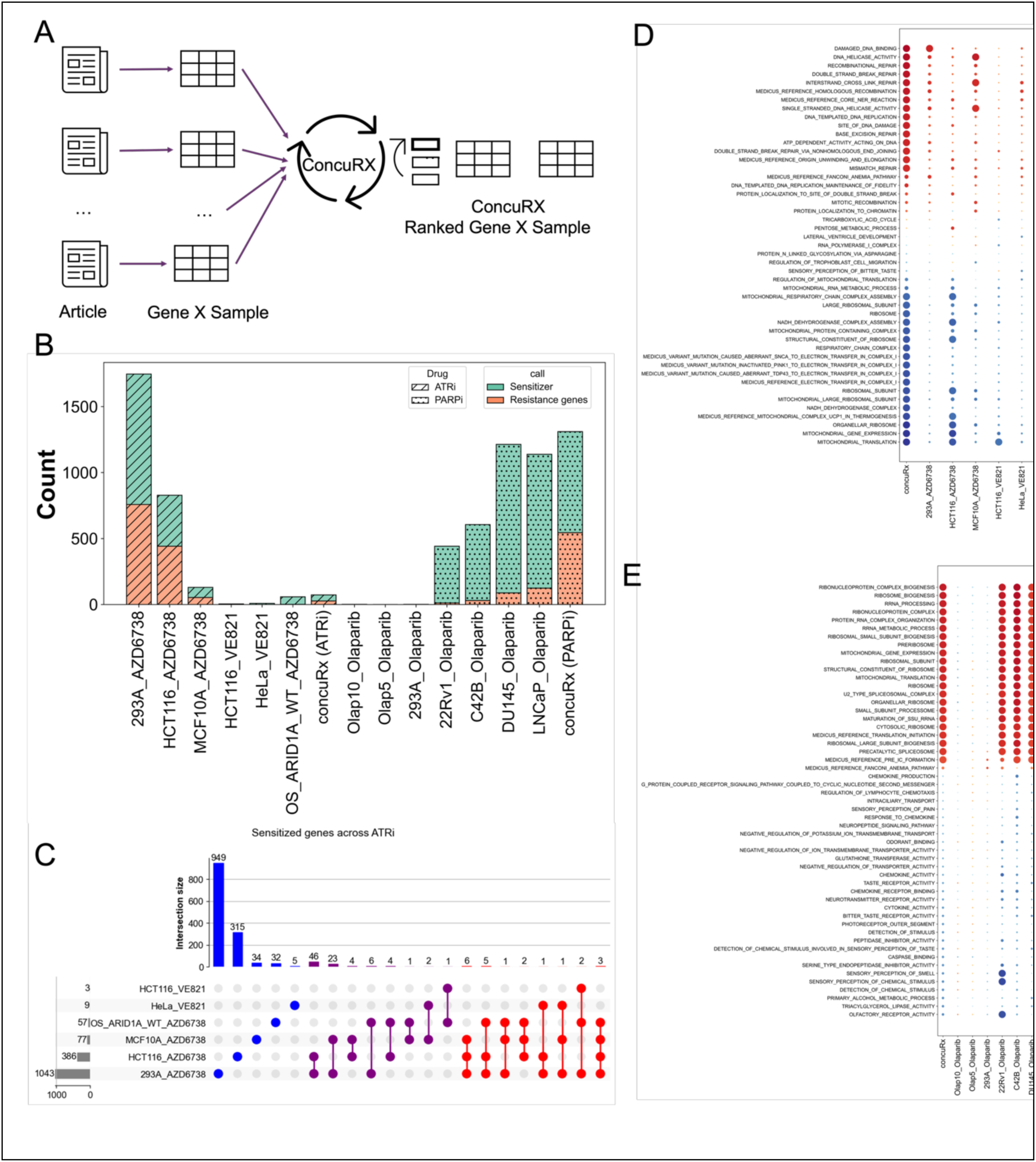
Workflow and Data Summary. A) A) ConcuRx workflow; B) Number of Sensitized and Resistance genes across individual datasets and concuRX; C) Overlapping methods applied to the ATRi sensitized genes across datasets; D-E) GSEA results from ATRi and PARPi CRISPR data.

### CRISPR-screen datasets

We curated genome-wide CRISPR knockout screens from three ATR inhibitor (ATRi) and three PARP inhibitor (PARPi) studies (Supplementary Table S1). The ATRi collection comprised six screens: AZD6738/ceralasertib screens in 293A, HCT116, and MCF10A cells ^20^; an AZD6738 screen in U2-OS ARID1A-proficient cells ^21^; and VE821 screens in HCT116 and TP53-deficient HeLa cells ^22^. The PARPi collection comprised seven olaparib/AZD2281 screens: A2780 ovarian carcinoma cells ^23^; four prostate cancer cell lines (LNCaP, C4-2B, 22Rv1, and DU145) representing different TP53 backgrounds ^24^; and 293A cells ^25^. Dataset sources, treatments, cell lines, processing methods, and gene-level statistics are summarized in Supplementary Table S1.

For each screen, we obtained either author-processed gene-level results or raw read-count data that were reprocessed locally using the corresponding analysis workflow. MAGeCK-derived datasets provided gene-level ranking statistics, log-fold changes, and left- and right-sided *P*-values ^26^, whereas drugZ-derived datasets provided normalized Z-scores (NormZ) and synthetic-lethal/sensitization and suppressor/resistance *P*-values (pval_synth and pval_supp) ^27^. For each drug–cell line comparison, we retained a gene-level effect statistic and its associated significance measure for subsequent harmonization and integration.

### Data ingestion and harmonization

Gene-level results from individual CRISPR screens were harmonized by identifying the corresponding gene-symbol, effect-size, and significance fields for each dataset. Gene symbols were standardized for capitalization and leading or trailing whitespace before datasets were merged. Because the source studies reported significance measures and effect statistics in different formats, dataset-specific transformations were applied to obtain comparable gene-level inputs for integration. When separate one-sided *P*-values were provided for effects in opposite directions, they were converted to a two-sided *P*-value as *P*_two-sided_ = min(1, 2 min(*P*_left_, *P*_right_)). P-values reported on logarithmic scales were back-transformed according to metadata-defined transformations. Values reported as (log_10_(*P*) ) were transformed as 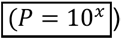, whereas values reported as (−log_10_(*P*)) were transformed as 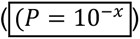 where *x* denotes the reported value. Because gene-level effect statistics differed in scale among source-analysis pipelines, effect values were standardized within each screen when required: 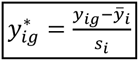 where *y_ig_* is the reported effect for gene (g) in screen (i), and (*y̅_i_*) and (*s_i_*) are the mean and standard deviation of gene-level effects in screen (i), respectively. Dataset-specific transformations were applied consistently to all genes within each screen. The harmonized datasets were then merged using the union of observed gene symbols, with each screen contributing one standardized effect estimate and one two-sided *P*-value for each available gene. Genes not evaluated or reported in each screen were retained as missing observations and were not imputed.

### Gene-level random-effects meta-analysis

Random-effects meta-analysis was performed independently for each gene using the metafor R package ^28^. A gene was included in the meta-analysis only when an effect estimate and corresponding *P*-value were available from at least five individual screens. Genes represented in fewer than five evaluable screens were retained with missing meta-analysis statistics and were not imputed.

Let *y_ig_* denote the harmonized effect estimate for gene *g* in screen *i*, and *P_ig_* its corresponding two-sided *P*-value. Because source datasets did not uniformly provide standard errors, the standard error for each effect estimate was derived from its two-sided *P*-value. *P*-values smaller than 10^−16^ were set to 10^−16^, and Standard errors were bounded below by 10^−16^.

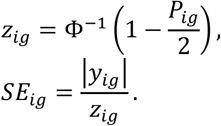

For each gene, a random-effects model was fitted using the *rma* function in metafor, with the Paule–Mandel estimator for between-study variance (method = "PM") and the Knapp–Hartung adjustment for inference (test = "knha"). The model was defined as

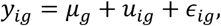

where *μ_g_* represents the mean cross-study effect, *u_ig_* represents between-study variation with variance 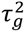, and *ε_ig_* represents within-study sampling error with variance 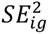.

The resulting integrated random-effects estimate, *μ*_RE_, its standard error and confidence interval, associated (*P*_RE_), estimated between-study variance 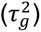, heterogeneity statistics (I^2^), and number of contributing screens (*k*) were retained for downstream analyses.

For sensitivity analysis, the same gene-level models were fitted using the DerSimonian–Laird estimator (method = "DL"), with the same derived standard errors and other model settings unchanged.

### Sensitization and Resistance classification

Negative and positive ConcuRx integrated effect estimates ( *μ*_RE_ ) represented sensitization and resistance, respectively, consistent with the directional interpretation of effect estimates across the individual input screens. Effect-size thresholds for classifying genes as sensitizers or resistance-associated genes were determined using the breakpoint-estimation procedure described below. The same breakpoint procedure was applied independently to the ConcuRx integrated effect distribution and to each individual dataset.

### Breakpoint estimation

To define sensitization and resistance thresholds from the continuous gene-level effect distributions, we applied a two-segment piecewise linear regression separately to the positive and negative effect tails. For resistance, positive effect sizes were ordered from largest to smallest; for sensitization, negative effect sizes were ordered from most to least negative. We evaluated all candidate breakpoints while requiring a minimum of 20 genes in each segment. For each candidate breakpoint *k*, separate linear regressions were fitted to the two segments, and the breakpoint minimizing the total residual sum of squares (RSS) was selected:

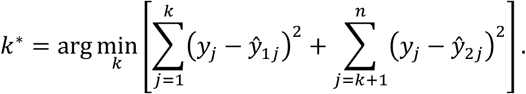

The effect size immediately preceding the selected breakpoint was used as the corresponding sensitization or resistance threshold.

### Quantification of missed sensitization and resistance calls

For each literature-curated gene, we counted the number of individual datasets in which the gene exceeded the sensitization breakpoint ( *c_S_* ) and the number in which it exceeded the resistance breakpoint (*c_R_*). Genes with (*c_S_* > *c_R_*) and a concordant negative ConcuRx effect were classified as expected sensitizers, whereas genes with (*c_R_* > *c_S_*) and a concordant positive ConcuRx effect were classified as expected resistance genes. Genes with equal numbers of sensitizing and resistance calls (*c_S_* = *c_R_* = 0) were not assigned a direction and were excluded from this analysis.

For an expected sensitizer, an individual dataset was considered to have missed the gene when the gene did not satisfy the dataset-specific sensitization criterion. Analogously, an expected resistance gene was considered missed when it did not satisfy the resistance criterion. The fraction of missed calls was calculated across evaluable gene–dataset comparisons separately for sensitization and resistance.

### Rank-improvement analysis

For each literature-curated gene, ConcuRx and individual-screen ranks were compared to determine whether cross-study integration improved prioritization in the biologically expected direction. To account for differences in the number of genes represented across datasets, raw ranks were converted to normalized ranks:

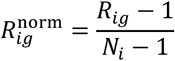

where *R_ig_* is the raw rank of gene *g* in dataset *i*, and *N_i_* is the total number of ranked genes in that dataset. Normalized ranks therefore ranged from 0 (top of the ranking; strongest sensitization) to 1 (bottom of the ranking; strongest resistance).

For sensitization-associated genes, normalized rank improvement was calculated as

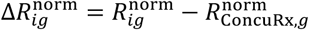

whereas for resistance-associated genes, the direction was reversed:

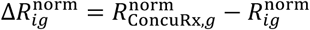

Thus, positive values indicated improved prioritization by ConcuRx in the biologically expected direction for both sensitization- and resistance-associated genes. Rank improvement was summarized across gene–dataset comparisons for the literature-curated modules. For each literature-defined module, the standard deviation of normalized gene ranks was calculated for ConcuRx and for each individual dataset. The individual-dataset values were summarized by their median for each module. Differences in within-module rank dispersion between ConcuRx and the individual datasets were then evaluated across modules using a Wilcoxon signed-rank test, pairing the ConcuRx spread with the median individual-dataset spread for the corresponding module.

### Pathway-level ranking coherence

Pathway-level coherence was evaluated using the literature-defined sensitization and resistance modules listed in Table 1. For each module, we calculated the standard deviation of the ranks of its constituent genes and, separately, the standard deviation of their effect sizes in ConcuRx and in each individual dataset. Lower within-module dispersion indicated that genes belonging to the same literature-defined module occupied more similar positions in the ranking or exhibited more similar effect estimates.

**Table 1:** Summary of hypothesis supported by individual, overlapped genes and concuRx.

| Drug | Hypothesis | Description | Genes Involved |
| --- | --- | --- | --- |
| ATRi | H1 | POLE3/POLE4 polymerase epsilon accessory complex are ATRi sensitizers | POLE3, POLE4 |
|  | H2 | Loss of RNASEH2 complex sensitizes cells to ATRi | RNASEH2A, RNASEH2B, RNASEH2C |
|  | H3 | The ATR checkpoint/replication-stress response pathway is a significantly enriched ATRi-sensitization module | ATR, ATRIP, CHEK1, ETAA1, CLSPN, RAD17, RAD9A, RAD1, HUS1, RHNO1 |
|  | H4 | The HR/Fanconi anemia repair module gene set overlaps with pathways previously linked to ATR inhibitor sensitivity. | BRCA1, BRCA2, PALB2, BRIP1, MAD2L2, FANCA, FANCB, FANCC, FANCD2, FANCG, FANCM, FAAP24, FAAP100, RAD50, RAD51D, XRCC2 |
|  | H5 | DNA replication-associated ATRi sensitization module | MCM9, DSCC1, FEN1, MCM2, RPA2, MCM6, RPA1, RFC2 |
|  | H6 | The PARP/BER/SSB repair module is one of the most consistently sensitizing groups to ATR inhibition | PARP1, XRCC1, APEX1, APEX2, ERCC1 |
|  | H7 | PP2A-associated genes form a high-confidence ATRi sensitization cluster | LCMT1, PPP2R4, PPP2R1A, PPP2R5D, KIAA1524/CIP2A |
|  | H8 | KLF5/chromatin-associated transcription–replication stress module | KLF5, BRD4, SMARCA4, ARID1B, CNOT1, SETD1A, THRAP3, KIAA1524/CIP2A |
|  | H9 | Cell-cycle and Mediator-associated genes define a resistance-leaning ATRi response group. | MED12, CCNC, CDK8, CDC25A, CDC25B, CDK2, CCNA2 |
|  | H10 | Oxidative/metabolic stress-associated genes are resistant to ATRi | EEF1B2, LUC7L3, KNTC1, SOD2, LIAS, RETSAT, KEAP1, NAE1 |
| PARPi | H11 | 16 kinase gene module | CLK1, PI4KA, PRKX, PLK1, MET, CDK5, CAMK1, ROCK2, JAK1, JAK2, LIMK2, ULK2, HCK, IRAK1, TBK1, TNIK |
|  | H12 | Loss of RNASEH2 complex sensitizes cells to PARPi | RNASEH2A, RNASEH2B, RNASEH2C, TOP1 |
|  | H13 | C12orf5/TIGAR as an olaparib-sensitizing gene | C12orf5/TIGAR, BRCA1, BRCA2, RAD51, CCNE2, BLM, CDC45, MCM10, MKI67, PCNA |
|  | H14 | DDR-associated PARPi sensitizers and CHEK2-mediated resistance | MMS22L, ESCO2, RNASEH2B, WDR76, MCM6, POLB, CHD1, FANCI, RAD54L, HELLS |
|  | H15 | CRPC-associated PARPi resistance genes | PARP1, ARH3, UBR5, YWHAE, USP48, TEAD4, CCND1 |

The frequency with which ConcuRx produced lower within-module rank or effect-size dispersion than individual datasets were summarized across pathway–dataset comparisons.

### Gene-set enrichment analysis

Pre-ranked gene-set enrichment analysis (GSEA) was performed using the prerank function in GSEApy^29^. Gene rankings were generated using the ConcuRx integrated effect estimates *μ*_RE_ or the corresponding effect estimates from individual CRISPR screens. Gene sets were obtained from MSigDB v2025.1.Hs, including the C5, C2.CP.KEGG_MEDICUS, C2.CP.REACTOME, and C2.CP.WIKIPATHWAYS collections, comprising 19,554 gene sets in total. GSEA was performed with a minimum gene-set size of 3, maximum gene-set size of 1,000, 1,000 permutations, and a random seed of 6. The same gene-set collections and analysis parameters were applied to ConcuRx and individual-screen rankings to enable direct comparison of pathway-level enrichment.

### Perturbation-based stability analysis

Robustness of gene prioritization to input perturbation was evaluated for the six ATRi screens by adding random noise to one screen at a time while leaving the remaining five screens unchanged. For each perturbed screen, independent additive noise was sampled from a normal distribution with mean 0 and standard deviation 1 and added to the gene-level effect values. The perturbation procedure was repeated 200 times for each of the six screens, resulting in a total of 1,200 perturbed ATRi dataset collections.

For each perturbation, ConcuRx meta-analysis and simple averaging were repeated using the same procedures applied to the unperturbed data. For the overlap-based approach, sensitizer genes were identified independently in each individual screen using the corresponding breakpoint-derived threshold, and the intersection of the sensitizer sets across all six screens was defined as the overlap-based gene set. The same procedure was repeated after each perturbation.

Stability of the selected sensitizer gene sets was quantified using Jaccard similarity between the sets obtained from the original and perturbed data:

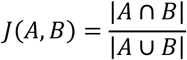

where A and B represent the sensitizer sets obtained from the original and perturbed data, respectively. Jaccard similarity was calculated independently for ConcuRx, simple averaging, and the overlap-based approach across all 1,200 perturbation replicates.

Stability of the continuous gene ranking was assessed using Kendall’s Tau between rankings derived from the original and perturbed data. For each comparison, rankings were restricted to genes present in both the original and perturbed datasets so that the two rankings contained the same set of genes. Kendall’s Tau was calculated for ConcuRx and the average-based approach using scipy.stats.kendalltau. The overlap-based approach was not included in the rank-correlation analysis because it produces a selected gene set rather than a continuous genome-wide ranking.

### Software and reproducibility

ConcuRx preprocessing, harmonization, comparative analyses, enrichment analyses, and robustness analyses were implemented in Python, and gene-level random-effects meta-analysis was performed in R. The code and data are provided in GitHub repo.

### Statistical Analyses

All quantification and statistical analysis were done using established R and Python packages, published tools, or in house codes. The details are provided in the individual Methods sections above.

### Ethics Approval

Not Applicable

## RESULTS

### ConcurX workflow to prioritize drug MOA genes

ConcuRx integrates gene-level effects across heterogeneous datasets to prioritize reproducible drug mechanism-of-action (MOA)-associated genes while accounting for uncertainty and between-study variability. The framework takes as input gene-level effect sizes and associated *P*-values obtained from standard screen-processing pipelines, such as MAGeCK^26^ or drugZ^27^ for CRISPR-screen data. For each gene, ConcuRx estimates an integrated effect across datasets while modelling between-study heterogeneity using a random-effects meta-analysis framework. Between-study variance is estimated using the Paule–Mandel estimator (Methods; Figure 1A), and genes are subsequently ranked according to their integrated effects and statistical support.

To evaluate ConcuRx, we curated 13 published genome-wide CRISPR-KO screens, comprising six ATR inhibitor (ATRi) and seven PARP inhibitor (PARPi) datasets from six studies and ten cell lines (Table S1). Across the six ATRi and seven PARPi screens, respectively, 21,466 and 21,363 unique genes were represented, of which 16,658 and 15,280 had estimable random-effects effect sizes (μ_RE) and associated *P*-values (*P*_RE) in ConcuRx (Methods; Table S2). To define sensitization and resistance calls from the continuous ConcuRx rankings, we applied segmented linear regression separately to the sensitizing and resistance distributions to identify effect-size breakpoints (Methods). Combining these breakpoints with statistical significance identified 46 sensitizers and 57 resistance genes for ATRi, and 765 sensitizers and 545 resistance genes for PARPi (Figure 1B).

The individual screens differed substantially in their identification of sensitization and resistance genes. The original studies used study-specific thresholds for effect size and statistical significance to identify sensitization and resistance hits (Table S1; Figures 1B and S1). Consequently, the number and identity of significant genes varied substantially among screens, and requiring overlap across multiple datasets progressively reduced the number of retained genes (Figure 1C; Figure S2). Simple averaging, in contrast, incorporates effect magnitude but does not account for differences in statistical uncertainty or between-study heterogeneity.

As an initial assessment of biological relevance, we performed pre-ranked gene-set enrichment analysis (GSEA) using ConcuRx rankings and compared the resulting pathway signals with those obtained from individual screens (Methods). For ATRi, sensitization-associated genes were enriched for DNA replication and DNA-damage repair, whereas resistance-associated genes were enriched for mitochondrial translation and ribosomal processes^20,22^ (Figure 1D). For PARPi, sensitization-associated signals included ribosome biogenesis, rRNA processing, ribonucleoprotein-complex assembly, translation, and mitochondrial gene expression, whereas resistance-associated signals included chemokine/GPCR signaling, ion transport, sensory receptor activity, and other cellular processes. Individual CRISPR screens showed more fragmented and context-dependent enrichment patterns, whereas ConcuRx produced more coherent enrichment across functionally related processes, suggesting that cross-study integration can recover pathway-level signals that are inconsistently represented in individual screens^24,30^ (Figure 1E).

We next systematically evaluated whether ConcuRx could recover biologically relevant genes missed by conventional approaches, capture pathway-level signals, improve prioritization of MOA-associated genes under cross-study variability, and recover mechanisms supported by independent evidence. Because a comprehensive ground-truth set of ATRi- and PARPi-response genes is unavailable, we curated 15 experimentally supported hypotheses from 13 published studies, including studies whose datasets were not incorporated into ConcuRx (Table 1). These hypotheses provided the biological reference framework for the subsequent evaluation of ConcuRx.

### ConcurX recovers biologically relevant genes missed by standard approaches

To determine whether ConcuRx recovers biologically relevant genes obscured by study-specific noise, we evaluated the literature-curated hypotheses described above (Table 1), focusing here on H1 and H2. According to the H1, POLE3 and POLE4, accessory subunits of DNA polymerase ε, are strong ATR-inhibitor sensitizers^20–22,31^. Hustedt et al. experimentally showed that loss of POLE3 or POLE4 markedly sensitizes cells to ATR inhibition^22^. Their experiments further indicated that this sensitivity was not explained by the histone-deposition function of the POLE3/POLE4 complex; the authors instead proposed that perturbation of Pol ε activity or another replisome-associated function may underlie the dependency on ATR-mediated replication-stress protection. In ConcuRx, this is one of the strongest two-gene modules: POLE3 ranks 1 with an effect size of −4.84, and POLE4 ranks 4 with an effect size of −3.86. In the individual datasets, one out of six datasets (HCT116_VE821) missed calling POLE3 as sensitizer (Figure 2A-B). While POLE4 is negative in all six datasets, it falls below the sensitizer breakpoint in the rank plot of the same HCT116_VE821 dataset (Figure 2A-B). In this example, the POLE3/4 would have been missed by any overlap-based method while ConcurRX correctly retains it as one of the highest-ranked ATRi sensitization genes (Figure 2C).

**Figure 2.**
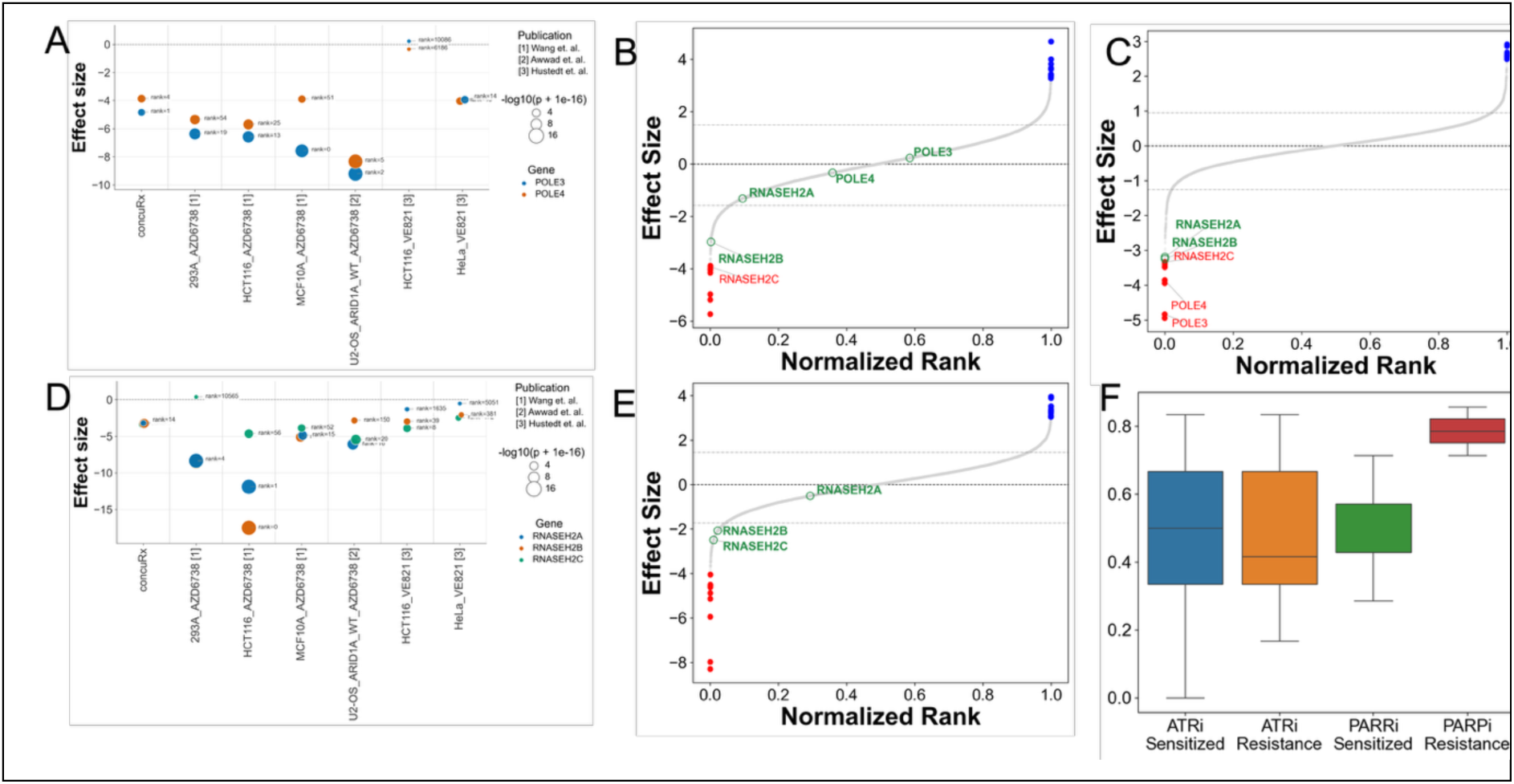
Noise Reduction. A) POLE3/4 (H1) module genes. B-C) Effect sizes sorted by normalized ranks by HCT116_VE821 and ConcuRx respectively showing the H1 model gene; D) RNASEH2 (H2) module genes; E) Effect sizes sorted by normalized ranks by HeLa_VE821 showing the H2 model gene; F) Fractions of times a call is missed but recovered by ConcuRX.

The next experimentally validated hypothesis, H2 states the RNASEH2 complex as one of the clearest and most reproducible ATRi synthetic-lethal modules. RNASEH2A, RNASEH2B, and RNASEH2C form a defined enzymatic complex involved in removing ribonucleotides incorporated into genomic DNA and processing RNA/DNA hybrids. Wang et al. identified RNASEH2 deficiency as a major determinant of ATRi sensitivity and experimentally showed that loss of RNASEH2A or RNASEH2B sensitizes cells to AZD6738 and VE822 in vitro, with RNASEH2B loss also sensitizing xenograft tumors to AZD6738 in vivo^20^ ^22^. This dependency was highly reproducible across studies: the Hustedt/Durocher consensus analysis identified all three RNASEH2 subunits among only 11 genes recovered in at least four of seven ATRi screens^22^. Consistent with these findings, Awwad et al. recovered RNASEH2A/C, together with other established ATRi-response genes such as ATM and POLE3/POLE4, in genome-wide CRISPR screens comparing ARID1A-WT and ARID1A-deficient cells^21^. Together, these studies support RNASEH2 deficiency as a recurrent ATRi vulnerability linked to replication-associated DNA damage rather than a broader, nonspecific R-loop response. In ConcuRx, RNASEH2C, RNASEH2B, and RNASEH2A rank 8, 12, and 13, with effect sizes of −3.34, −3.25, and −3.19, respectively. This reproduces the major conclusion of Wang et al. ^20^, where RNASEH2 deficiency was the prominent ATRi synthetic-lethal dependency and was validated experimentally in vitro and in xenografts. The results from ConcuRx show why integration matters: RNASEH2A is negative in all six individual datasets, yet RNASEH2A falls below the sensitizer breakpoint in 2 datasets: Hela and HCT116 from Hustedt et. al., and RNASEH2C is slightly positive in 293A cell line from Wang et al.^20^ [Figure 2B, 2D-E, S3]. A strict intersection approach could penalize RNASEH2A/C because of these discordant datasets, but ConcuRx preserves the correct biological conclusion that loss of the RNASEH2 complex sensitizes cells to ATR inhibition (Figure 2C). Wang et al. observed RNASEH2C as being sensitized in wet lab, but their CRISPR data was not supporting it suggesting it as a potential screen-to-screen variation^20^.

Similar to the two hypotheses described above, we extended this analysis to the hand-curated hypothesis/gene modules in Table 1 and evaluated how often their associated genes were missed by individual datasets but recovered by the ConcuRX ranking (Methods). A gene was classified as a sensitizer (resistance) gene if it showed sensitizing (resistance) effects in more individual datasets than effects in the opposite direction, with the overall direction concordant with the ConcuRX ranking. Within each dataset, an effect was called only when its magnitude exceeded the sensitizer or resistance breakpoint defined using the criteria from the corresponding source study. Across the curated modules, 54 genes were classified as sensitizers and 10 as resistance genes. Among the individual-dataset evaluations of these genes, 50% (43%) of the expected sensitizer calls and 42% (78%) of the expected resistance calls of ATRi (PARPi) were missed by the individual analyses but recovered in the correct direction by concuRX (Figure 2F). Collectively, these results show that concuRX recovers biologically supported signals that are frequently missed in individual CRISPR screens, supporting the use of meta-analysis to reduce study-specific noise and improve robust gene prioritization.

### ConcurX captures pathway-level signal more effectively

Genes that modulate drug response are functionally coordinated, such that sensitizing and resistance-associated genes converge on distinct biological pathways rather than occurring as isolated gene-level effects^32,33^. In previous section, we demonstrated via GSEA analysis that the ConcuRx gene ranking identifies the relevant pathways known in the literature in more coherent manner than the individual datasets. While ConcuRx captured these pathways, we next investigated whether genes within the literature-defined modules exhibited greater coherence in their integrated effects and rankings. To investigate the pathway-level signal, we used 12 (3) pathways from the hypothesis Table 1 which had been experimentally demonstrated as sensitized (resistant) pathway. Among them we discuss 2 pathways (H3, H4) from the hypothesis table in the following, while H5 is presented in the Supplementary.

The ATR checkpoint/replication-stress response module (H3) includes ATR, ATRIP, CHEK1, ETAA1, CLSPN, RAD17, RAD9A, RAD1, HUS1, and RHNO1. Wang et al. identified ATR-pathway and replication-stress response genes among ATRi co-essential genes across three genome-wide AZD6738 CRISPR screens^20^, supporting the broader principle that genetic disruption of ATR-dependent checkpoint signalling can increase sensitivity to ATR inhibition. This relationship was particularly evident in the Hustedt/Durocher consensus analysis, in which ATRIP was among only 11 genes recovered as ATRi-sensitization hits in at least four of seven screens, while ATR itself was identified as a sensitization hit in three screens^22^. Because ATRIP is an essential partner of ATR in checkpoint signalling, reduced ATRIP function is expected to decrease ATR-pathway activity and increase dependence on residual ATR function. Together, these findings support ATR checkpoint and replication-stress response genes as a coherent ATRi-sensitization module in which genetic weakening of ATR-dependent signalling can increase vulnerability to further pharmacological ATR inhibition. Consistent with these previous findings, ConcuRX recovered a strong and coherent sensitization signal across the H3 module, with nearly all genes exhibiting negative integrated effects. RHNO1, ATRIP, HUS1, and ATR were among the strongest sensitizers within the module (Figure 3A-B). Examination of the individual datasets, however, showed study-to-study variability: while RHNO1 and ATRIP showed sensitizing effects across all six datasets, HUS1 was sensitizing in five of six (Figure 3C), and other module genes were not consistently identified as sensitizers in every screen (Figure 3A). Thus, individual screens can miss otherwise recurrent sensitization signals, whereas ConcuRX integrates evidence across datasets to preserve the broader pathway-level pattern despite study-specific variability.

**Figure 3.**
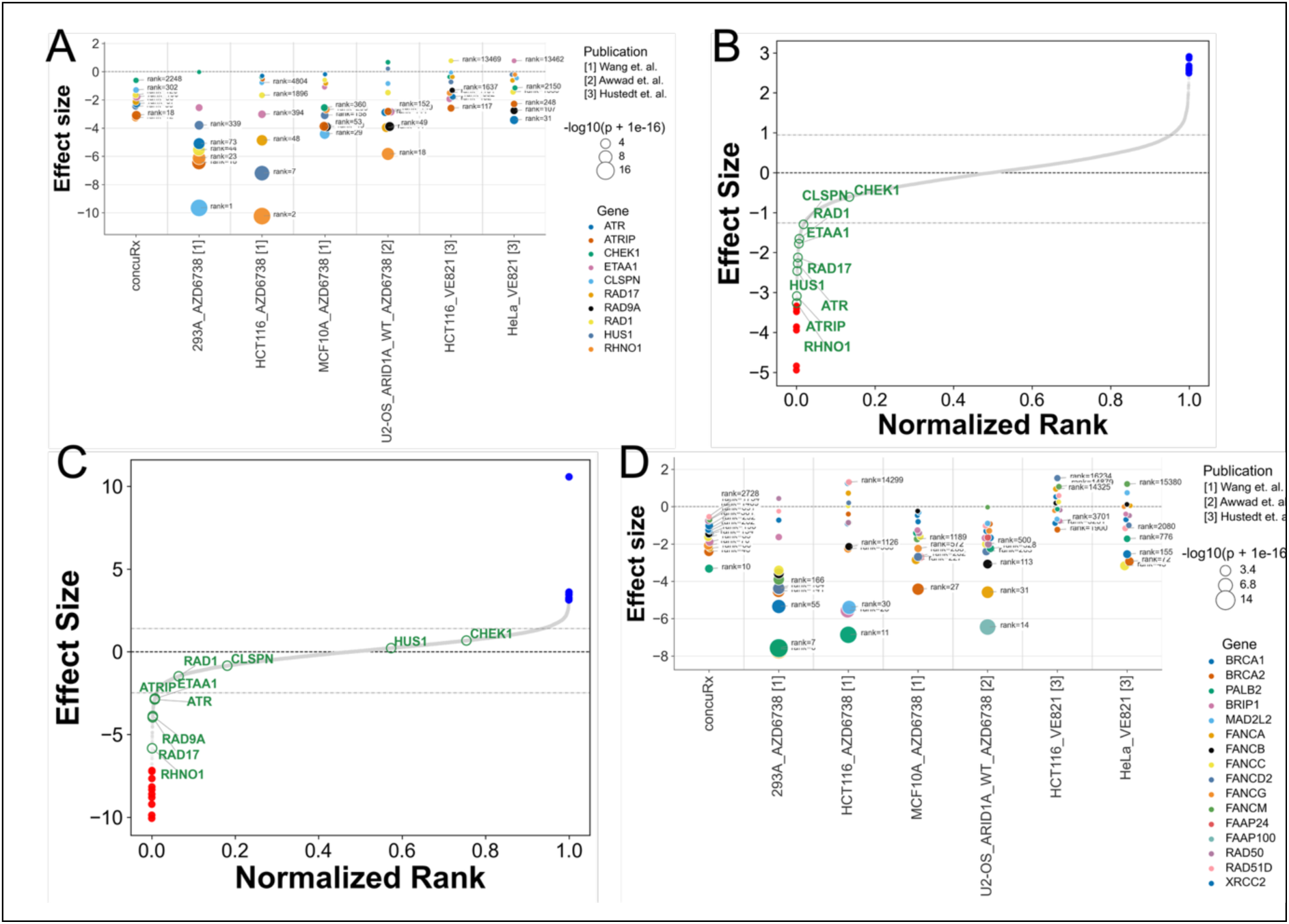
Pathway signal. A) ATR checkpoint/replication-stress response (H3) module genes; B-C) Effect sizes sorted by normalized ranks by ConcuRx and U2-OS_ARID1A_WT_AZD6738 dataset respectively showing the H3 model gene; D) HR/Fanconi anemia repair (H4) module genes.

Next, we examined H4, the HR/Fanconi anemia (FA) repair module, comprising BRCA1, BRCA2, PALB2, BRIP1, MAD2L2, FANCA, FANCB, FANCC, FANCD2, FANCG, FANCM, FAAP24, FAAP100, RAD50, RAD51D, and XRCC2. Multiple lines of prior evidence connect defects in HR/FA-mediated DNA repair with increased dependence on replication-stress response pathways. Wang et al. identified HR- and FA-related genes among ATRi-sensitization hits across genome-wide AZD6738 CRISPR screens^20^, consistent with broader evidence that BRCA/HR-deficient cells can become dependent on ATR–CHK1 signaling^34,35^. The biological coherence of this module is further supported by Cai et al., who showed that loss of FA/BRCA pathway genes sensitizes cells to ATM inhibition and demonstrated functional cooperation between ATM and FA/BRCA pathways in DNA double-strand-break end resection^36^. More broadly, large-scale genetic mapping of the DNA-damage response has also identified HR and Fanconi anemia genes as recurrent determinants of cellular responses to genotoxic stress^37^. Together, these studies support HR/FA deficiency as a biologically coherent source of DNA-repair and replication-stress vulnerability, providing a rationale for evaluating whether ConcuRX recovers this module as a shared ATRi-sensitization signal. Consistent with this prior evidence, ConcuRx recovered a strong sensitization signal across the H4 HR/Fanconi anemia module, with 14 of 16 genes showing negative integrated effects. PALB2, BRCA2, FANCA, FANCG, and BRIP1 were among the strongest sensitizers (Figure 3D). Several of these genes, including BRCA2, PALB2, BRIP1, and MAD2L2, were also identified as ATRi co-essential hits by Wang et al.^20^. The broader recovery of HR and Fanconi anemia genes in the integrated ranking extend these previously reported individual-gene observations and supports a coherent pathway-level ATRi sensitization signal.

Across the 15 pathways from the hypothesis list in Table 1, we computed the spread (standard deviation) of gene ranks and effect sizes in ConcuRx and compared these with the corresponding spread in individual datasets (Methods; Figure S5). For 86% (67%) of the pathway–dataset comparisons, the spread of gene ranks (effect sizes) was reduced in ConcuRx, indicating greater within-module coherence in the integrated ranking and effect estimates. Consistent with this observation, the within-module rank spread was significantly lower in ConcuRx than across individual datasets (Wilcoxon signed-rank test, median P = 0.0029296875 across ConcuRx vs individual datasets). Although all genes within a module are not necessarily expected to exhibit identical sensitizing or resistance effects, their closer positioning in the ConcuRx ranking indicates that genes belonging to the same literature-defined module are more consistently prioritized together^38^.

### ConcurX improves ranking of drug MOA genes in the presence of cross-study variability

As the genes within a drug-relevant pathway tend to exhibit coordinated sensitizing or resistance effects, we hypothesized that a good ranking method would improve their collective prioritization. We therefore assessed whether genes belonging to sensitizing (resistance) pathways were ranked more strongly toward the sensitizing (resistance) end by ConcuRX, compared with their rankings in individual datasets. To evaluate the rank improvement, we focused on H6 and H7 hypothesis in detail, while H8 is presented in the Supplementary (Figure S6).

The PARP/BER/SSB repair module includes PARP1, XRCC1, APEX1, APEX2, and ERCC1, which function across interconnected pathways involved in the recognition and repair of single-strand and base-associated DNA lesions. APEX2 provides particularly strong prior evidence linking this module to ATRi sensitivity: Hustedt et al. identified APEX2 among only 11 genes recovered as ATRi-sensitization hits in at least four of seven screens and experimentally validated its contribution to ATRi sensitivity^22^. Mechanistically, APE2 has been implicated in ATR-dependent checkpoint signalling from single-strand breaks^39^ and is also synthetic lethal with BRCA1/BRCA2 deficiency^40^, further linking single-strand-break processing to replication-stress vulnerabilities. More broadly, Olivieri et al. identified BER/SSB-repair genes as determinants of cellular responses across a large collection of DNA-damaging perturbations^37^. Consistent with this biological evidence, ConcuRx recovered a strong sensitization signal across the PARP/BER/SSB repair module, with all five genes showing negative integrated effects and PARP1, APEX2, XRCC1, and APEX1 among the highest-ranked sensitizers (Figure 4A-B). Importantly, examination of the individual ATRi screens illustrates the study-to-study variability that ConcuRx is designed to address: PARP1 and APEX1/APEX2 showed sensitizing effects across all six datasets, whereas XRCC1 showed the same direction in five of six (Figure 4C). Despite this variability, ConcuRx recovered a coherent module-level sensitization pattern and substantially improved the sensitization ranks of these genes relative to the individual datasets: improved by mean normalized ranking of 0.05 (corresponding to a 5-percentage-point shift toward the sensitizing end of the genome-wide ranking) and in more than 83% of the datasets (Figure 4D-E). Together, these results show that ConcuRx preserves recurrent DNA-repair vulnerabilities and improves their prioritization while accounting for study-specific experimental and biological heterogeneity.

**Figure 4.**
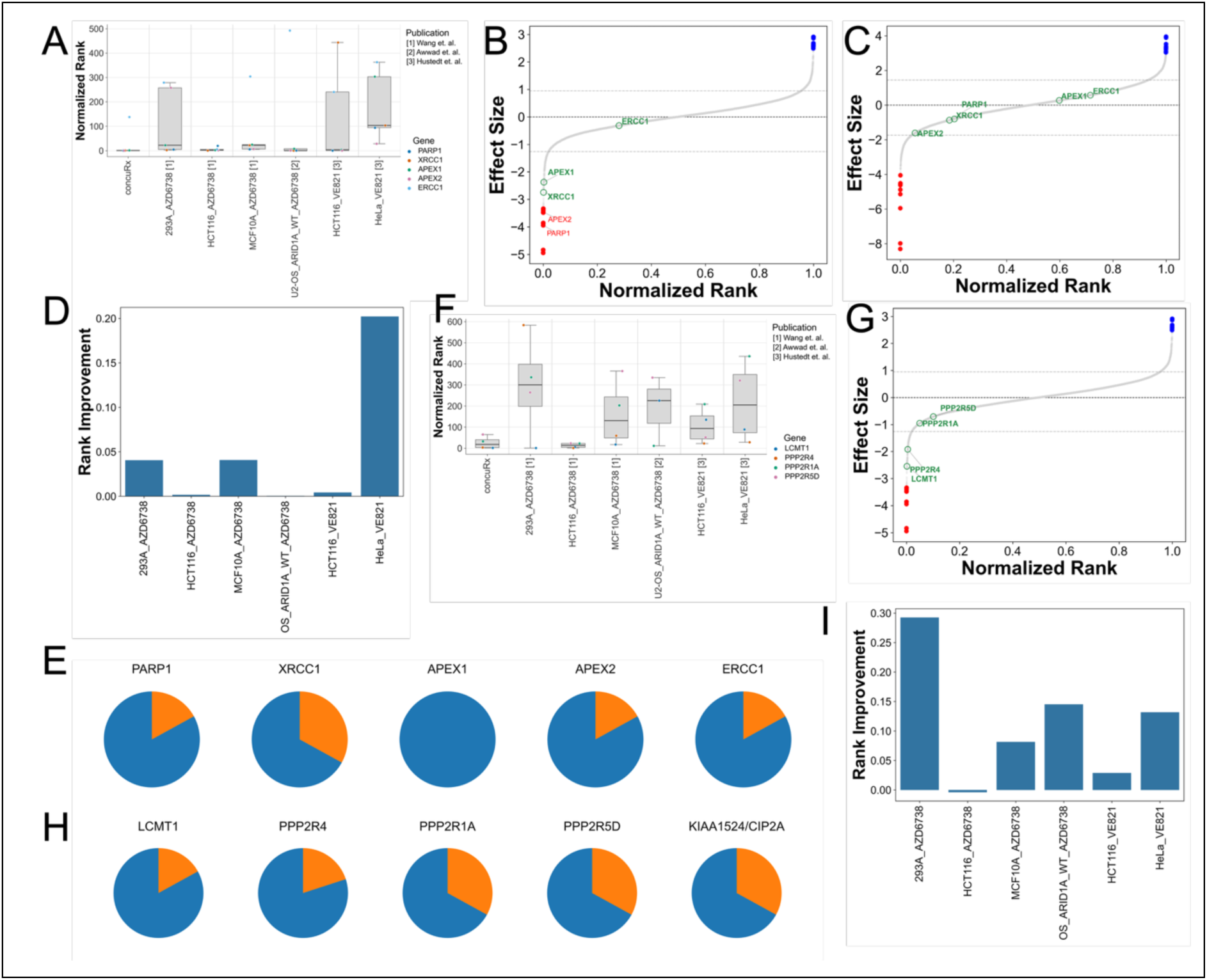
Rank Improvement. A) Normalized Rank distributions in PARP/BER/SSB (H6) module; B-C) Effect sizes sorted by normalized ranks by ConcuRx and HeLa_VE821 dataset respectively showing the H6 model genes; D) Rank improvement in the H6 module; E) Percentage of datasets showing rank improvement for the H6 model genes, Blue (orange) denotes improved (not improved); F) Normalized Rank distributions in PP2A-associated genes (H7) across datasets; G) Effect sizes sorted by normalized ranks by ConcuRx showing the H7 genes; H) Percentage of datasets showing rank improvement for the H7 model genes; and I) Rank improvement in the H7 module.

The PP2A-associated module, comprising LCMT1, PPP2R4, PPP2R1A, PPP2R5D, and KIAA1524/CIP2A, showed substantial sensitization-rank improvement in the ConcuRx analysis, with improvement observed across a majority of the datasets. Prior studies provide support for several components of this module. Wang et al. identified LCMT1 among ATRi-sensitization candidates and noted that its role in an ATR-dependent or parallel pathway remained unresolved ^20^. Hustedt/Durocher et al. subsequently identified KIAA1524/CIP2A as one of only 11 genes recovered as an ATRi-sensitization hit in at least four of seven screens, supporting its role as a recurrent ATRi vulnerability^22^. In ConcuRx, the module showed an overall shift toward ATRi sensitization, although the signal varied across genes (Figure 4F-G). KIAA1524/CIP2A was the strongest member, ranking 6th overall with an integrated effect of −3.40 and showing a negative effect in all six paper datasets. LCMT1 also showed strong support, ranking 27th with an integrated effect of −2.55 and negative effects across all six datasets. PPP2R4 showed a negative ConcuRx effect (−1.92; rank 83) but greater variability, including one positive signal in the Wang et al. 293A dataset. The structural/regulatory subunits PPP2R1A and PPP2R5D were also shifted toward sensitization but ranked substantially lower: across datasets, the reduced mean normalized ranking is 0.11 and in more than 66% of the datasets (Figure 4H-I). Together, these results suggest that ConcuRx recovers a PP2A-associated ATRi-sensitization pattern driven most strongly by KIAA1524/CIP2A and LCMT1, while providing weaker support for other PP2A components. This is consistent with the recurrent identification of KIAA1524/CIP2A across independent ATRi screens^22^.

Across all the genes spanning 15 modules from the hypothesis list of Table 1, we quantified the rank improvements by comparing ConcuRx rankings with those from the individual datasets (Methods). ConcuRx improved prioritization in 71% of the gene-dataset comparisons, reflected by lower sensitization ranks for sensitized genes or higher resistance prioritization for resistance genes.

### ConcurX recovers independently supported mechanisms using studies not included in the integration

The hypotheses discussed above were derived from studies whose datasets were included in the ConcuRx analysis. Next, we evaluated hypotheses H9-11 derived from independent studies whose datasets were not integrated by ConcuRx, providing an independent validation of the integrated results. H9 and H10 are presented here, while H11 is described in the Supplementary (Figure S7).

We next examined H9: an ATRi resistance-associated module comprising MED12, CCNC, CDK8, CDC25A, CDC25B, CDK2, and CCNA2. Several members have independent experimental support as determinants of ATRi resistance. Schleicher et al. demonstrated that MED12 loss promotes resistance to multiple ATR inhibitors through activation of TGFβ signaling and restoration of replication-fork progression during ATR inhibition^41^. Interestingly, depletion of other Mediator or CDK8-module components, including MED7, MED13, CDK8, and CCNC, did not reproduce this phenotype in their experimental system. In contrast, Lloyd et al. independently showed that loss of Cyclin C/CCNC or CDK8 promotes ATRi resistance by reducing DNA hybrids and transcription-associated replication stress^31^, suggesting that the contribution of the CDK8 module can depend on cellular or experimental context. CDC25A provides another independently validated resistance mechanism: loss of CDC25A confers ATRi resistance by preventing the premature mitotic entry induced by ATR inhibition^42^. Together, these studies establish MED12, CCNC/CDK8, and CDC25A as mechanistically supported but distinct routes to ATRi resistance. Notably, the datasets from these mechanistic studies ^31,41,42^ were not included in the data integrated by ConcuRx. Nevertheless, the resistance-associated direction was independently recovered from the publication datasets used in our analysis: all seven genes showed positive integrated effects in ConcuRx, with MED12, CCNC, CDC25A, CDC25B, and CDK2 among the strongest resistance-ranked genes, while CDK8 and CCNA2 showed weaker but concordant positive effects (Figure 5A). The individual datasets also generally showed resistance-like effects except Hela_VE821, in addition to the varied strength of the signals across experimental contexts (Figure 5B). Thus, ConcuRx recovers a resistance-associated gene set that is concordant with mechanisms established in independent studies not included in the integration (Figure 5C), while also capturing context-dependent variation among individual genes. This provides external support for the ability of ConcuRx to identify reproducible resistance signals across heterogeneous CRISPR screens.

**Figure 5.**
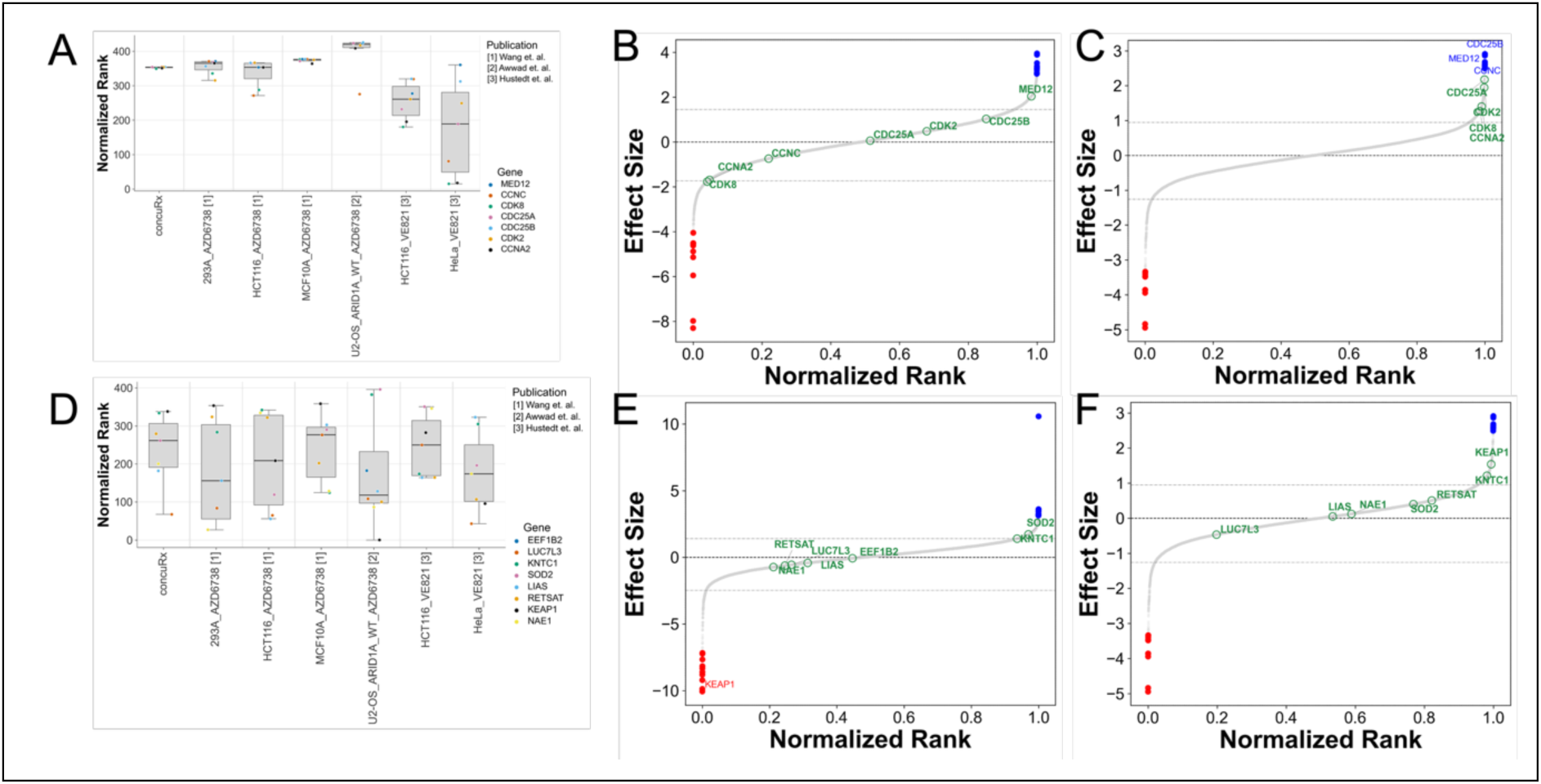
Independent Validation. Distributions of Normalized Ranks in ATRi resistance associate module: H9 (A) and H10 (D), B-C) Effect sizes sorted by Normalized Rank of Hela_VE821 and ConcuRx showing genes from H9 module, E-F) Effect sizes sorted by Normalized Rank of OS_ARIDA1A_WT_AZD6738 and ConcuRx showing H10 module genes. Blue and red dots are indicating top 10 resistance and sensitized genes; (See Figure S12 for H11 genes).

Hypothesis H10 examines a group of ATRi resistance-associated genes, including EEF1B2, LUC7L3, KNTC1, SOD2, LIAS, RETSAT, KEAP1, and NAE1. Schleicher et al. identified EEF1B2, LUC7L3, KNTC1, SOD2, LIAS, and RETSAT among the strongest common hits from high-dose VE822 and AZD6738 resistance screens and experimentally confirmed that their depletion increased ATRi resistance^41^. These genes act through diverse mechanisms: for example, LIAS depletion restored replication-fork progression during ATR inhibition, whereas LUC7L3 depletion shifted BCL-X splicing toward the anti-apoptotic isoform and reduced ATRi-induced apoptosis. Because the Schleicher datasets were not included in ConcuRx, recovery of these resistance-associated genes provides an independent test of the integrated analysis. ConcuRx recovered an overall positive integrated effect for this group (Figure 5D), consistent with resistance or reduced ATRi response and independently supporting the Schleicher findings. Other genes showed greater context dependence. NAE1 was selected for validation by Hustedt et al., but its ATRi-specific effect remained inconclusive because its depletion was highly cytotoxic even without ATR inhibition^22^. KEAP1 was associated with ATRi resistance by Wang et al.^20^, whereas the Awwad et al. screen showed sensitization in a different experimental context^21^. ConcuRx assigned KEAP1 a net positive effect, illustrating how cross-study integration can retain the predominant response while reducing the influence of a strong context-specific signal (Figure 5E-F).

Together, the independent evaluations in H9–H11 support the robustness of ConcuRx, which recovered previously reported sensitization and resistance patterns despite not inclusion of the studies from which these hypotheses were derived. These results demonstrate the value of cross-study integration for identifying reproducible drug-response signals across heterogeneous experimental contexts.

### ConcurX improves stability of gene ranking across datasets

We assessed ranking robustness to experimental noise by perturbing one sample at a time with additive noise, while retaining all samples, and recalculating gene rankings across repeated perturbations. Focusing primarily on ATRi sensitization, we evaluated robustness using two complementary metrics: Jaccard similarity, which measures preservation of the selected sensitizer/resistance gene sets, and Kendall’s tau, which measures agreement between rankings obtained from the original and perturbed data. For Jaccard similarity, ConcuRx was compared with average- and overlap-based approaches; for Kendall’s tau, ConcuRx was compared only with the average-based approach because the overlap method does not generate a continuous gene ranking.

ConcuRx showed substantially greater stability under perturbation. Jaccard similarities for ConcuRx remained above 0.6, compared with values below 0.3 for the average-based method, indicating greater preservation of sensitizer/resistance gene sets following noise addition (median pvalue of 7.2e-35 across datasets using one-sided Wilcoxon signed-rank test) (Figure 6A). The overlap-based method showed Jaccard similarities of closed to zero; most likely driven by very small, selected gene sets (approximately 3–5 top sensitizers; Figure 6B). Consistent with these results, Kendall’s tau correlations between original and perturbed rankings were consistently higher for ConcuRx than for the average-based approach (Figures 6C-D), demonstrating greater stability of the overall ranking (median pvalue of 0.001235 and 7.5e-05 for correlation statistics and significance across datasets using one-sided Wilcoxon signed-rank test). Together, the perturbation experiments indicate that ConcuRx provides more stable gene prioritization in the presence of noise and cross-study variability than the conventional approaches evaluated here.

**Figure 6.**
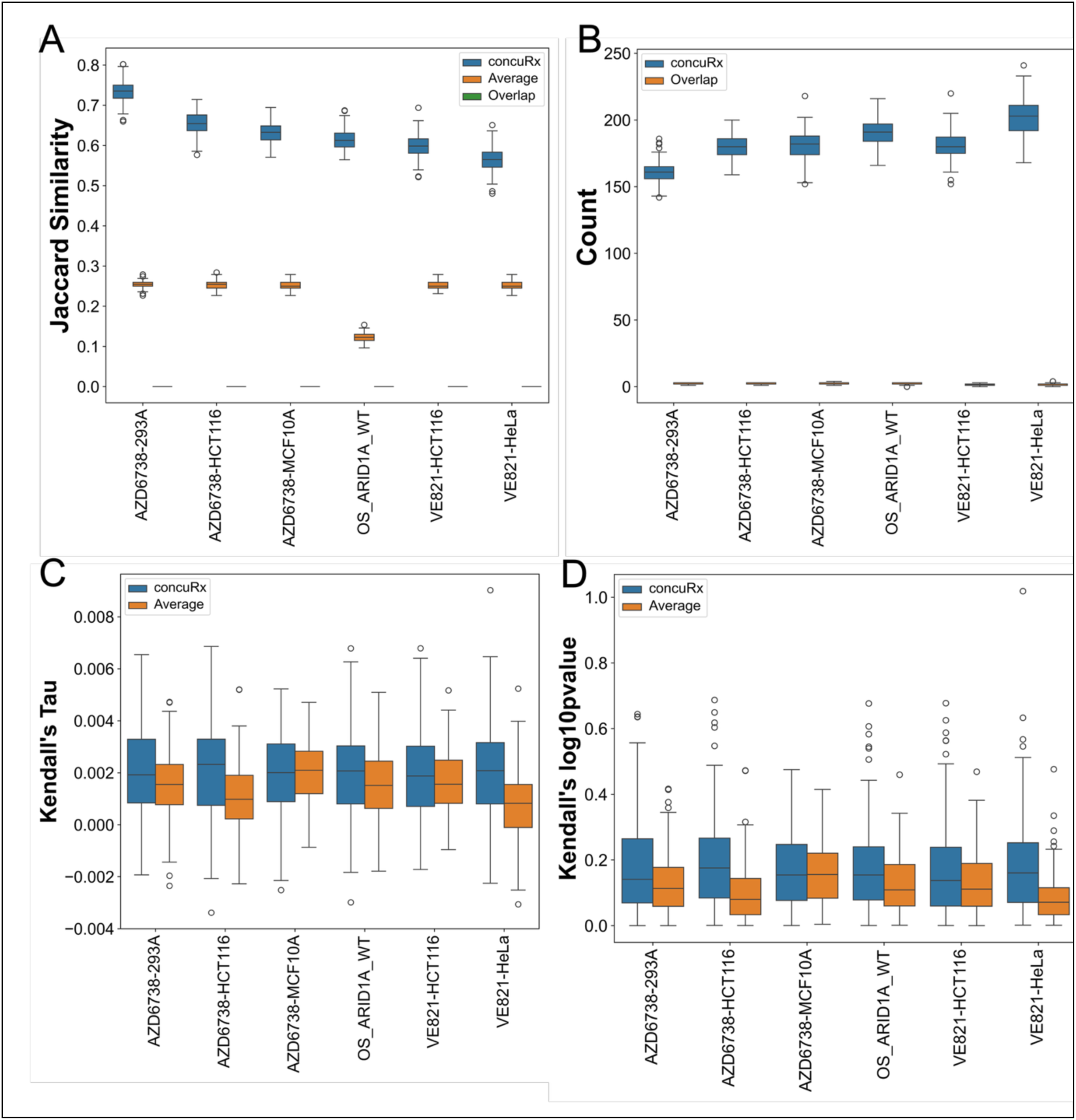
Stability. A) Jaccard Similarity of sensitized genes between actual data and noisy data obtained by ConcuRX (blue), Average (orange), Overlapped (green) method in ATRi datasets, B) Count of sensitized genes obtained by ConcuRx and overlap based approach, C) Kendall’s Tau of the actual and noisy data rankings obtained by concuRx and Average method, D) log10pvalue of Kendal’s rank correlation between actual and noisy data rankings obtained by concuRx and Average method.

We additionally assessed whether the integrated results were sensitive to the choice of random-effects procedure by comparing the PM-KnHa approach used in ConcuRx with the conventional DerSimonian–Laird (DL) approach. Effect estimates were broadly concordant between the two approaches, whereas differences were more apparent in the estimated between-study heterogeneity (Supplementary Results; Figure S12). Together, these analyses show that ConcuRx maintains more stable gene sets and rankings under data perturbation, while the comparison of random-effects procedures indicates that heterogeneity estimates can be sensitive to the choice of estimator when integrating a limited number of screens.

## DISCUSSION

We developed ConcuRx, a statistical framework for robust gene prioritization across heterogeneous CRISPR drug-response screens. When only a limited number of datasets are available and differences among cell lines, genetic backgrounds, treatment conditions, and experimental designs reflect biological as well as technical variation, conventional batch correction and pooled analysis may not be appropriate. ConcuRx instead retains study-level effects and integrates them through random-effects meta-analysis, explicitly modeling between-study variability rather than attempting to remove it. This provides a framework for identifying genetic dependencies that remain reproducible across heterogeneous experimental contexts while preserving information about study-specific variation.

Across ATRi and PARPi screens, ConcuRx recovered experimentally supported genes that could be missed by individual screens or strict overlap-based analyses. The POLE3/POLE4 and RNase H2 modules provide particularly clear examples. Both represent established ATRi vulnerabilities, yet individual datasets failed to identify a particular gene at the expected sensitization threshold. Notably, RNASEH2C showed weak or discordant evidence in an individual screen despite experimental support for RNase H2 deficiency as an ATRi vulnerability, whereas ConcuRx strongly prioritized all three RNase H2 subunits. These examples illustrate how requiring agreement among individually significant hits can discard biologically supported dependencies when otherwise recurrent signals are weakened in particular experimental contexts.

The benefit of integration also extended beyond individual genes to pathway-level biological signal. Literature-defined ATR checkpoint/replication-stress and HR/Fanconi anemia modules showed coherent sensitization patterns in the integrated ranking despite substantial variation among individual screens. Importantly, ConcuRx estimates each gene independently and does not use pathway membership when constructing the ranking. The increased coherence of genes belonging to experimentally supported modules therefore provides an independent indication that cross-study integration strengthens biologically structured signal rather than simply altering individual gene ranks. Across the literature-defined modules examined systematically, reduced within-module spread of ranks and effect sizes further supported greater pathway-level coherence after integration.

ConcuRx also improved the prioritization of MOA-associated genes under cross-study variability. The PARP/BER/SSB repair module, for example, showed substantial improvement toward the ATRi-sensitizing end of the integrated ranking relative to individual datasets. At the same time, integration did not uniformly promote every member of a predefined biological group. Within the PP2A-associated module, KIAA1524/CIP2A and LCMT1 received strong integrated support, whereas other components showed weaker or more variable effects. Thus, the improved prioritization observed for literature-defined modules reflects the accumulated evidence for individual genes rather than an imposed pathway-level constraint. Consistent with this interpretation, ConcuRx improved prioritization across a majority of gene–dataset comparisons and produced rankings that were more stable to data perturbation than average- and overlap-based approaches.

A further assessment came from hypotheses established by independent experimental studies whose datasets were not included in the ConcuRx integration. ConcuRx recovered resistance-associated signals consistent with independently characterized ATRi mechanisms, including genes associated with MED12-, CDK8/CCNC-, and CDC25-mediated resistance, despite variation in their effects among the datasets used for integration. Recovery of such mechanisms from a separate collection of screens provides additional evidence that ConcuRx can identify reproducible biological relationships that are not simply consequences of including the studies from which those hypotheses were originally derived.

These results also clarify how an integrated ConcuRx effect should be interpreted. Cross-study reproducibility is not equivalent to universal biological activity. A dependency experimentally validated in a particular cellular or genetic background may weaken or even reverse when examined across broader contexts. The Ipsen study provides a clear example: although PARP1, ARH3, UBR5, and YWHAE were experimentally validated as olaparib-resistance determinants in castration-resistant prostate cancer, ConcuRx reproduced the resistance direction for PARP1 and ARH3 but not UBR5 or YWHAE. This disagreement does not invalidate the context-specific experimental findings but suggests that some dependencies may be less reproducible across the broader cellular contexts represented in the integrated screens. Conversely, dependencies consistently supported across heterogeneous screens provide stronger evidence of cross-study reproducibility within the contexts examined. For example, the strong integrated sensitization of all three RNase H2 subunits, despite weaker or discordant signals in individual screens, illustrates a dependency that remains reproducible across experimental contexts. Thus, ConcuRx complements rather than replaces context-specific analyses: individual experiments establish dependencies within biological systems, whereas ConcuRx evaluates the extent to which those dependencies are supported across studies while retaining the underlying study-specific effects.

Comparison across drug classes suggests a broader application of this framework. ATR and PARP inhibitors exhibit synergistic activity, and their ConcuRx profiles showed convergence across multiple DNA-repair and replication-stress processes. The RNase H2 complex provided the clearest shared sensitization dependency, while additional signals involving HR/Fanconi anemia and replication-associated genes, including BRCA1/2 and MCM6, further supported convergence of the two drug-response landscapes. The analyses also identified resistance-associated DNA-repair dependencies, including CHEK2 and PARP1, indicating that shared genetic architecture may encompass both vulnerabilities and mechanisms that modulate treatment response. Independent experimental studies have linked RNase H2 deficiency to sensitivity to both drug classes and enhanced vulnerability to combined ATRi–PARPi treatment. Together, these observations illustrate how comparing integrated genetic-response profiles across drugs may identify shared pathways and complementary sensitization/resistance dependencies relevant to combination therapy. Such observations do not establish the mechanism of synergy by themselves but can generate evidence-grounded mechanistic hypotheses that prioritize pathways and genetic contexts for subsequent experimental testing.

Limitations and Future Directions

Several limitations remain. First, identification and harmonization of relevant CRISPR-screen datasets remain challenging, and the number of comparable screens available for many drugs is small. ConcuRx uses the Paule–Mandel estimator with Knapp–Hartung adjustment for random-effects inference in this setting; nevertheless, meta-analysis becomes increasingly uncertain as the number of available datasets decreases. For some genes or drugs, insufficient observations prevent reliable integrated estimation. In such settings, simpler approaches such as identifying recurrent overlapping hits may remain useful until additional datasets become available. Greater availability of standardized and well-annotated CRISPR drug-response datasets should therefore increase both the applicability and statistical power of cross-study integration.

A related limitation is that the integrated effect depends on the biological contexts represented among the available studies. If screens disproportionately represent a particular cancer type, genetic backgrounds, or experimental conditions, the resulting integrated effect should not be interpreted as universal across untested systems. Conversely, dependencies restricted to specific contexts may receive weaker integrated support despite being biologically valid within those settings. ConcuRx should therefore complement rather than replace context-specific analyses. As larger collections of screens become available, an important extension will be to stratify or explicitly model known sources of biological heterogeneity, enabling broadly recurrent dependencies to be distinguished from those associated with a particular molecular or cellular contexts.

ConcuRx, simple averaging, and alternative random-effects approaches may produce correlated effect estimates because they summarize the same underlying directional evidence. The advantage of ConcuRx therefore does not require fundamentally different pooled effects; rather, the framework incorporates study-level uncertainty and explicitly models between-study variability while providing statistical inference for gene prioritization. Among random-effects approaches, ConcuRx uses the Paule–Mandel estimator with Knapp–Hartung adjustment (PM-KnHa), rather than the conventional DerSimonian–Laird (DL) approach, because DL has known limitations when the number of studies is small^43,44^. Given the limited number of screens available in our analyses, we additionally compared PM-KnHa with DL to evaluate how the choice of random-effects procedure affects effect estimates and statistical uncertainty (Supplementary Results). Nevertheless, uncertainty and significance remain dependent on both the number and quality of contributing studies and should be interpreted together with the underlying study-level evidence.

Finally, although the present study evaluates ConcuRx using CRISPR drug-response screens, the underlying integration problem extends to other perturbational data types. Drug-treated proteomic, phosphoproteomic, transcriptomic, and other molecular-response studies similarly span different cell lines, compounds, doses, and experimental conditions, often with relatively few directly comparable datasets. Application of ConcuRx to these modalities will require appropriate definitions of effect size and uncertainty and systematic benchmarking against relevant biological evidence. Evaluating whether cross-study integration can recover reproducible molecular-response signals across these additional modalities represents an important direction for future work.

In summary, ConcuRx provides a framework for integrating heterogeneous functional-genomics screens while retaining information about cross-study variability. Across ATRi and PARPi datasets, ConcuRx recovered established drug-response genes and pathways, improved pathway coherence and gene prioritization, recovered mechanisms supported by independent studies, and produced more stable rankings under perturbation. Together, these findings support cross-study meta-analysis as a complementary strategy for distinguishing recurrent drug-response dependencies from context-dependent effects and provide a foundation for extending integrated genetic-dependency profiles toward mechanistic interpretation and drug-combination discovery.

## Supporting information

Supplementary Results

## ACKNOWLEDGEMENTS

The computational resources, in part, were provided by the NIH HPC Biowulf cluster.

## AUTHOR CONTRIBUTIONS

Conceptualization - MS; Data curation, Formal analysis, Software, Visualization – MS,MA;– MS,MA; Funding acquisition – EM; Investigation – MS; Methodology – MS,MA; Project administration – EM; Resources – EM; Supervision – MS; Validation – MA,EM; Writing – original draft - MS; Writing – review & editing – MS,EM.

The contributions of the NIH author(s) were made as part of their official duties as NIH federal employees, are in compliance with agency policy requirements, and are considered Works of the United States Government. However, the findings and conclusions presented in this paper are those of the author(s) and do not necessarily reflect the views or policies of the Department of Health and Human Services, nor does mention of trade names, commercial products, or organizations imply endorsement by the U.S. Government.

## SUPPLEMENTARY DATA

Supplementary Data will be available at NAR online.

## DATA AVAILABILITY

- This study did not generate new datasets.
- ConcuRx tool outputs are in the Supplementary data
- This paper analyses existing, publicly available data.
- The original code and the data analysis code is deposited at (https://github.com/mahfuzasharmin/concurx/). Any additional information required to reanalyze the data reported in this paper is available from the lead contact upon request.

## APPENDIX

Not applicable

