## Supplementary Results for "Robust Identification of Drug Mechanism of Action Associated Genes Across Heterogeneous Drug Treated Datasets"

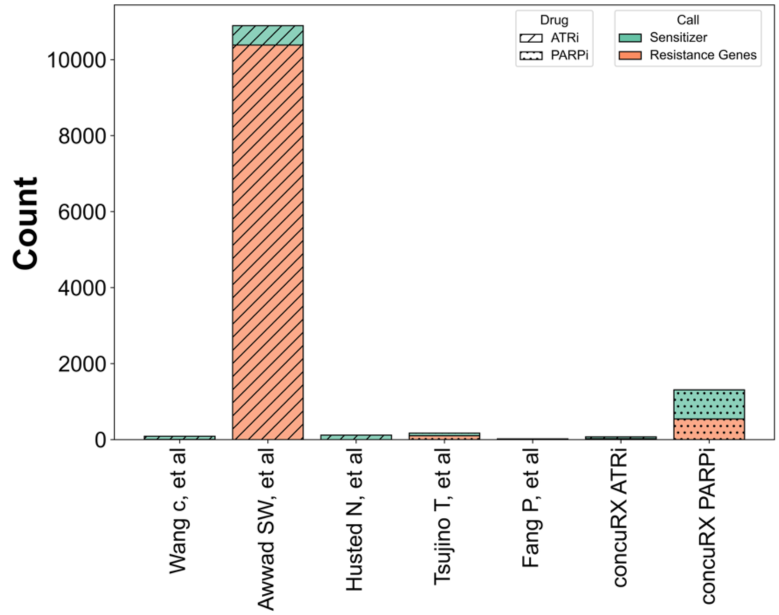


Figure S1: Number of sensitized and resistance genes obtained from the articles and ConcuRx

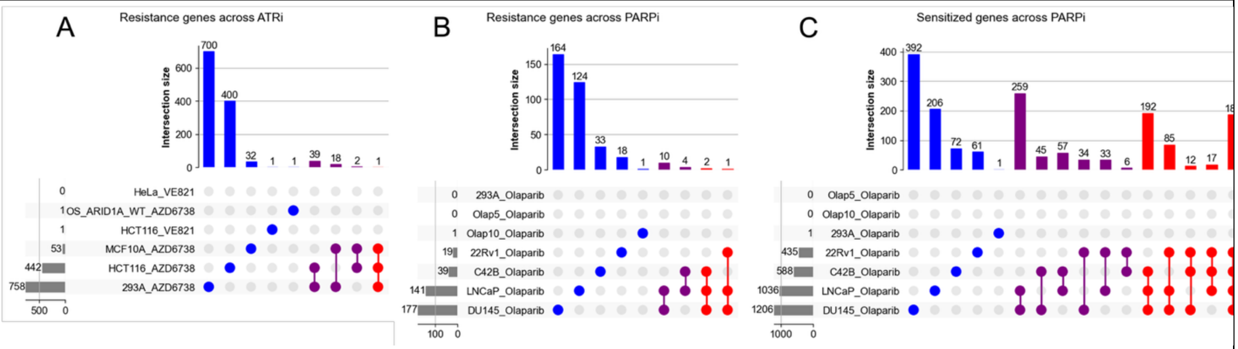


Figure S2: Number of genes shown using overlap-based method A) Resistance genes in ATRi datasets, B) Resistance genes in PARPi datasets, C) Sensitized genes in PAPRi datasets.

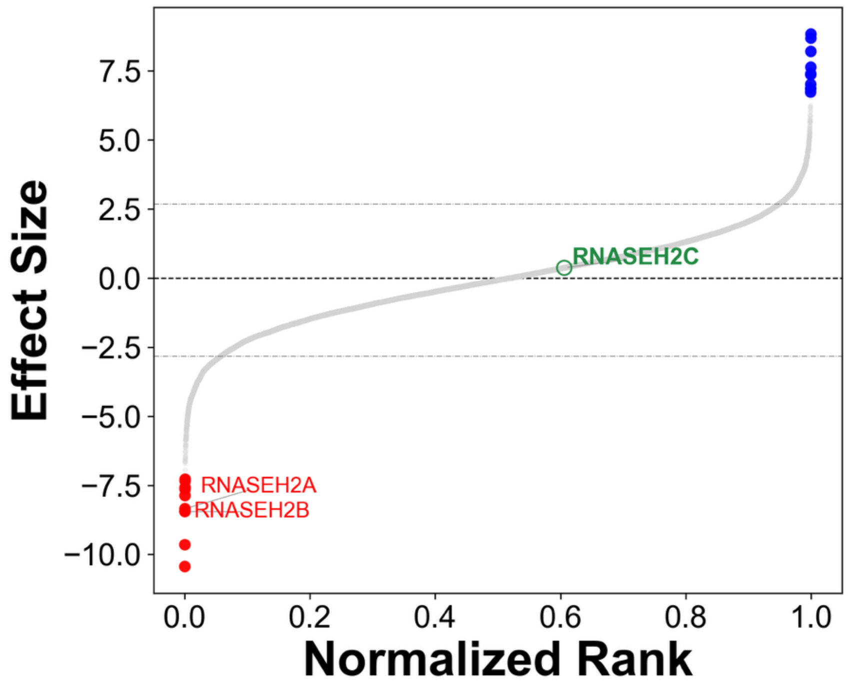


Figure S3: Effect sizes sorted by normalized ranks by 293A_AZD6738 showing the H2 model genes.

H5: Wang et al defines the broader DNA replication machinery module which includes MCM9, DSCC1, FEN1, MCM2, RPA2, MCM6, RPA1, RFC2, and related replication factors[1] . They identified DNA replication as a significantly enriched biological process among ATRi co-essential genes and highlighted replication-associated complexes, including the MCM and RFC complexes. Consistent with this finding, the ConcuRx results showed a broad shift toward sensitization across the module with MCM9, DSCC1, FEN1, MCM2, RPA2, and MCM6, among the highest-ranked sensitizers. Although many of these genes showed sensitizing effects across individual datasets, their effect sizes varied between screens. Thus, ConcuRx captures a coherent DNA-replication-associated sensitization signal across heterogeneous experimental contexts, supporting increased ATRi vulnerability when DNA replication functions are perturbed.

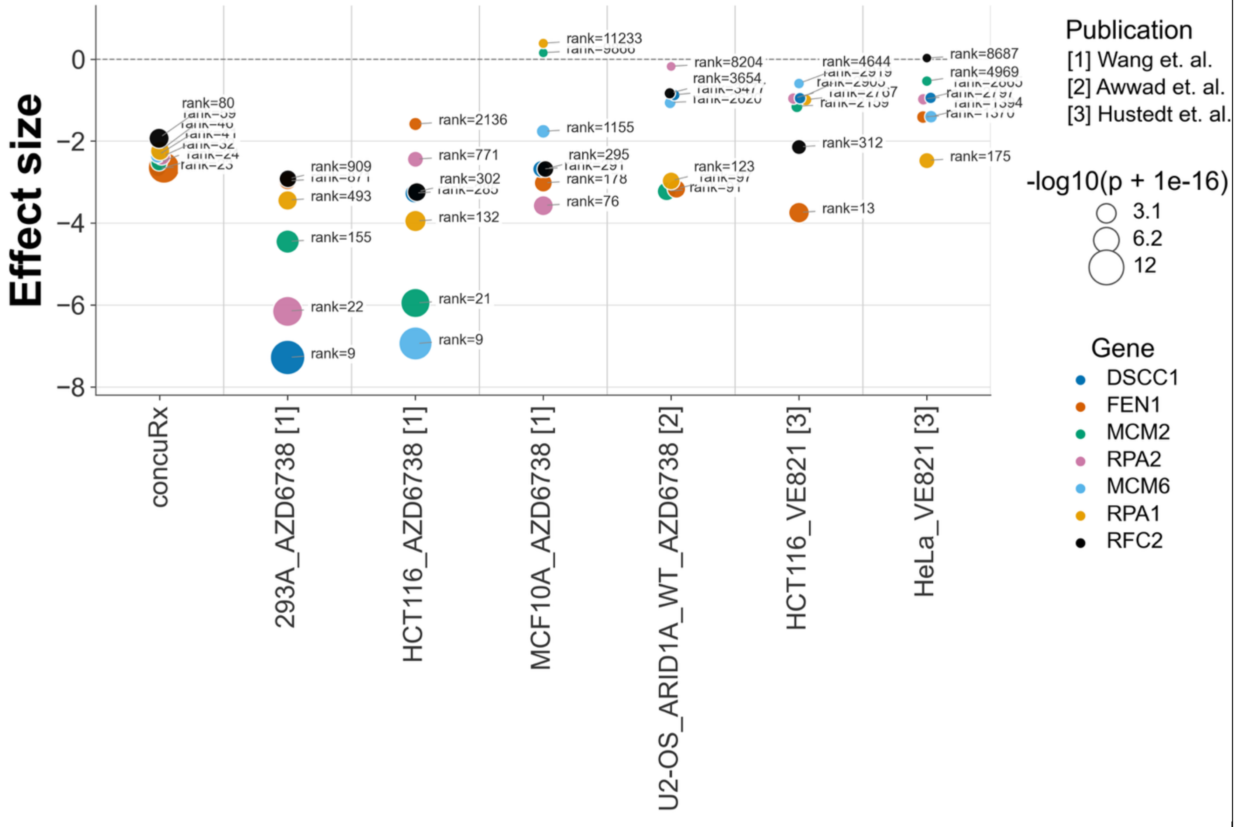


Figure S4: DNA replication machinery (H5) module genes.

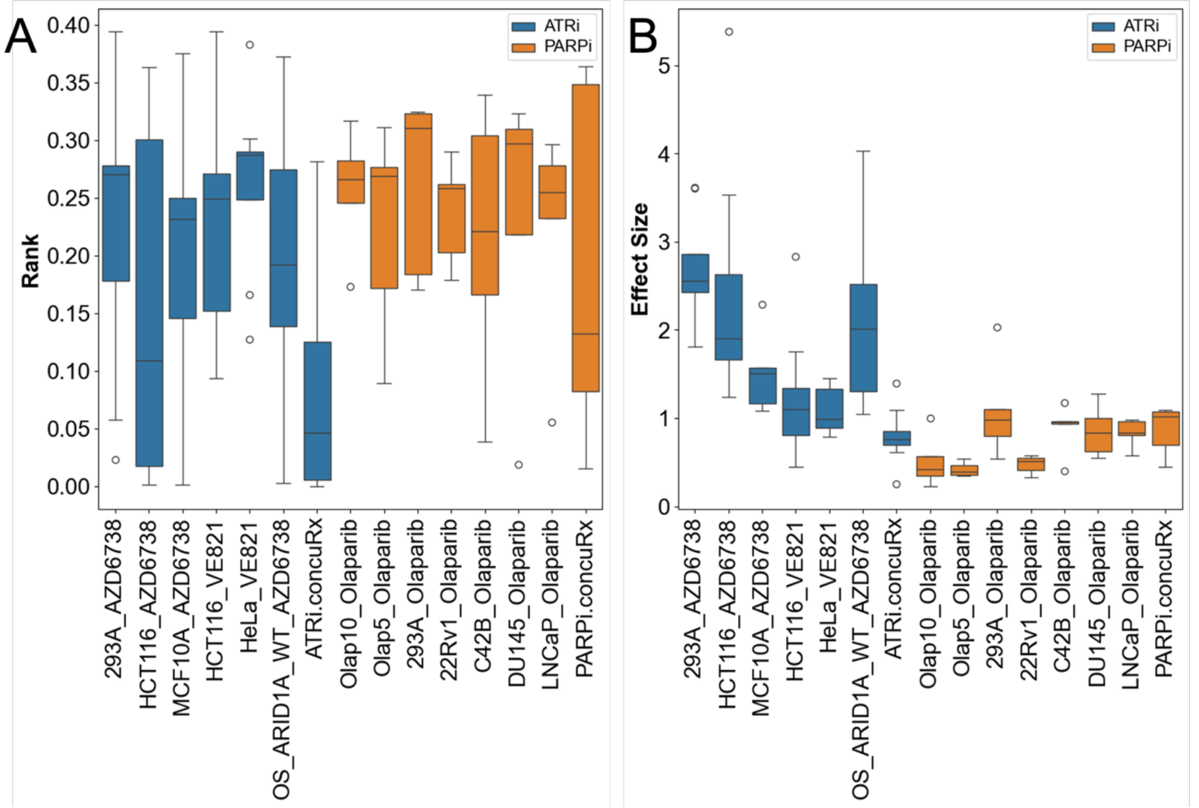


Figure S5: Spread (Standard deviation) of the Normalized ranks (A) and Effect sizes (B) across ConcuRx and individual datasets ranking.

H8: We additionally examined a curated KLF5/chromatin-associated transcription–replication stress module, comprising KLF5, BRD4, SMARCA4, ARID1B, CNOT1, SETD1A, THRAP3, and KIAA1524/CIP2A. This group showed substantial cross-study variability in ConcuRx. In ConcuRx, CNOT1, THRAP3, and SETD1A showed relatively strong sensitizing effects, whereas KLF5 showed a more moderate integrated effect. The KLF5 signal is consistent with Awwad et al., who experimentally demonstrated that KLF5 loss sensitizes cells to ATR inhibition and increases transcription–replication conflicts, DNA–RNA hybrids, replication stress, DNA damage, and genomic instability[2] . Mechanistically, KLF5 was shown to protect against transcription-associated replication stress, at least partly by regulating BRD4 recruitment to chromatin. In contrast, other members of the curated module were less consistent across studies; for example, SMARCA4 showed a strong sensitizing effect in the Awwad dataset but was near-neutral after cross-study integration. Thus, ConcuRx retains the experimentally supported KLF5-associated sensitization signal while moderating effects that are less reproducible across experimental contexts, illustrating how integration can reveal differences in the cross-study reproducibility of ATRi-associated vulnerabilities.

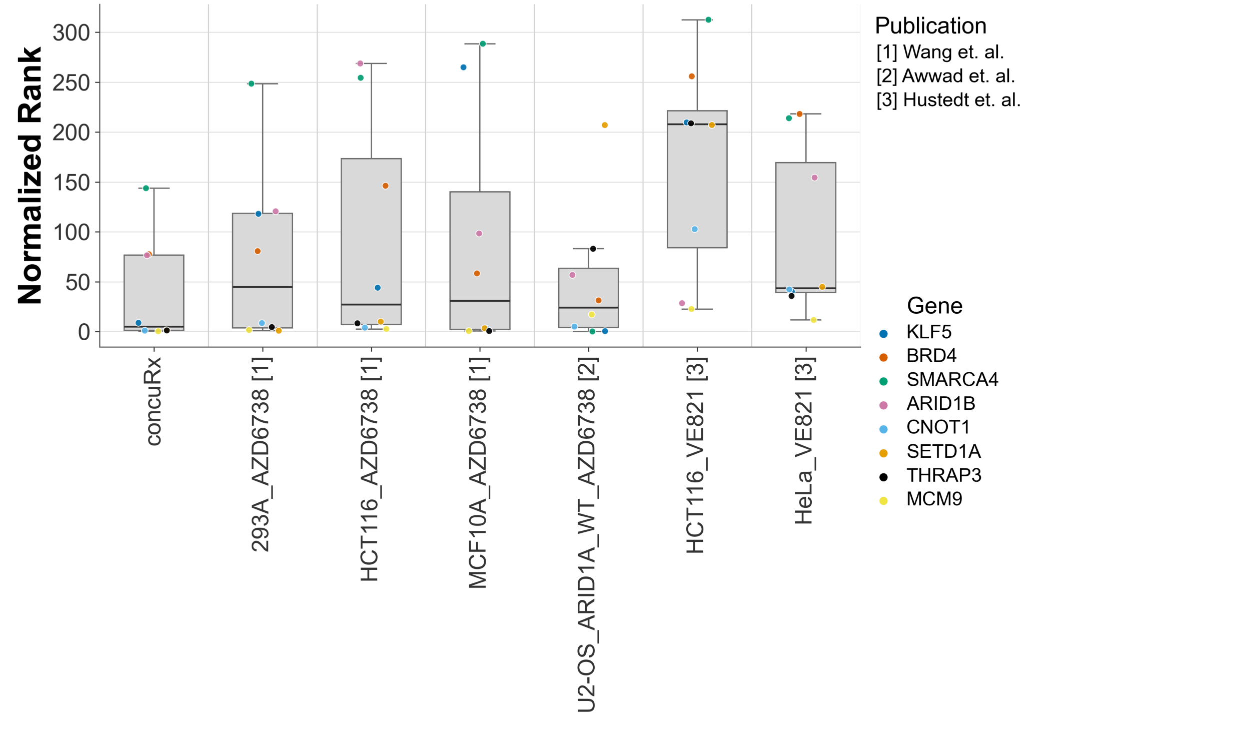


Figure S6: Normalized ranks of KLF5/chromatin-associated transcription–replication stress (H8) module genes.

H11: Liu et al. identified 16 candidate kinases (CLK1, PI4KA, PRKX, PLK1, MET, CDK5, CAMK1, ROCK2, JAK1, JAK2, LIMK2, ULK2, HCK, IRAK1, TBK1, and TNIK) associated with PARPi response, including PLK1, and subsequent drug screening highlighted CLK1 as their strongest synergistic target with olaparib. Because the Liu screen was not included in ConcuRx, we tested whether these candidates were supported across the seven independent olaparib screens. Among Liu's 16 candidates, PLK1 showed the strongest ConcuRx support, with a strong integrated sensitization effect and negative effects in six of seven datasets. PI4KA and CDK5 also showed consistent sensitizing directions across most or all individual screens, although their pooled evidence was weaker or unavailable. Thus, ConcuRx provided independent cross-study support for a subset of the kinase vulnerabilities identified by Liu et al., with PLK1 showing the strongest generalization across datasets.

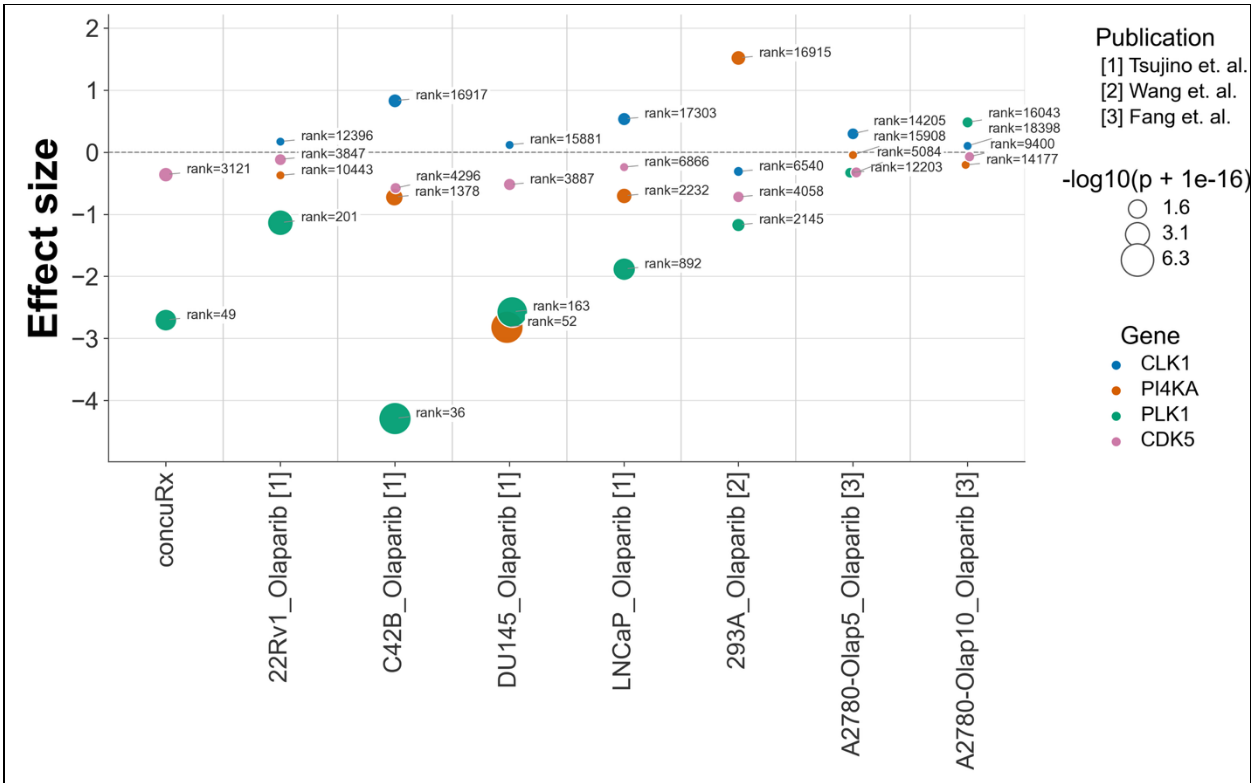


Figure S7: Effect size and significances of 16 candidate kinases (H11).

H12: Zimmermann et al.[3] identified RNase H2 deficiency as a strong PARPi vulnerability, showing that defective ribonucleotide excision repair causes accumulation of genomic ribonucleotides that are processed by TOP1 into lesions promoting PARP1 trapping, replication-associated DNA damage, and cell death. The authors demonstrated hypersensitivity of RNase H2-deficient cells to PARP inhibitors, including olaparib and talazoparib, and highlighted recurrent RNASEH2B deletion in subsets of cancers as a potentially exploitable vulnerability. ConcuRx independently recovered this mechanism across heterogeneous olaparib screens: all three RNase H2 subunits showed significant integrated sensitizing effects despite substantial variation in effect strength and significance across individual datasets. TOP1, a mechanistic component of the RNase H2/PARP-trapping model, was also recovered as a significant sensitizer despite weaker and dose-dependent effects in some individual screens.

This finding is particularly relevant because PARP and ATR inhibitors show synergistic activity, and RNase H2 deficiency has independently been associated with sensitivity to both drug classes and enhanced vulnerability to combined ATRi–PARPi treatment. Thus, recurrence of the RNase H2 complex across the ATRi and PARPi ConcuRx analyses provides a shared genetic vulnerability consistent with the biological basis of this combination. More broadly, this example suggests that comparing robust ConcuRx dependencies across drugs may help identify shared vulnerabilities and generate mechanistic hypotheses for drug combinations, which can subsequently be tested experimentally.

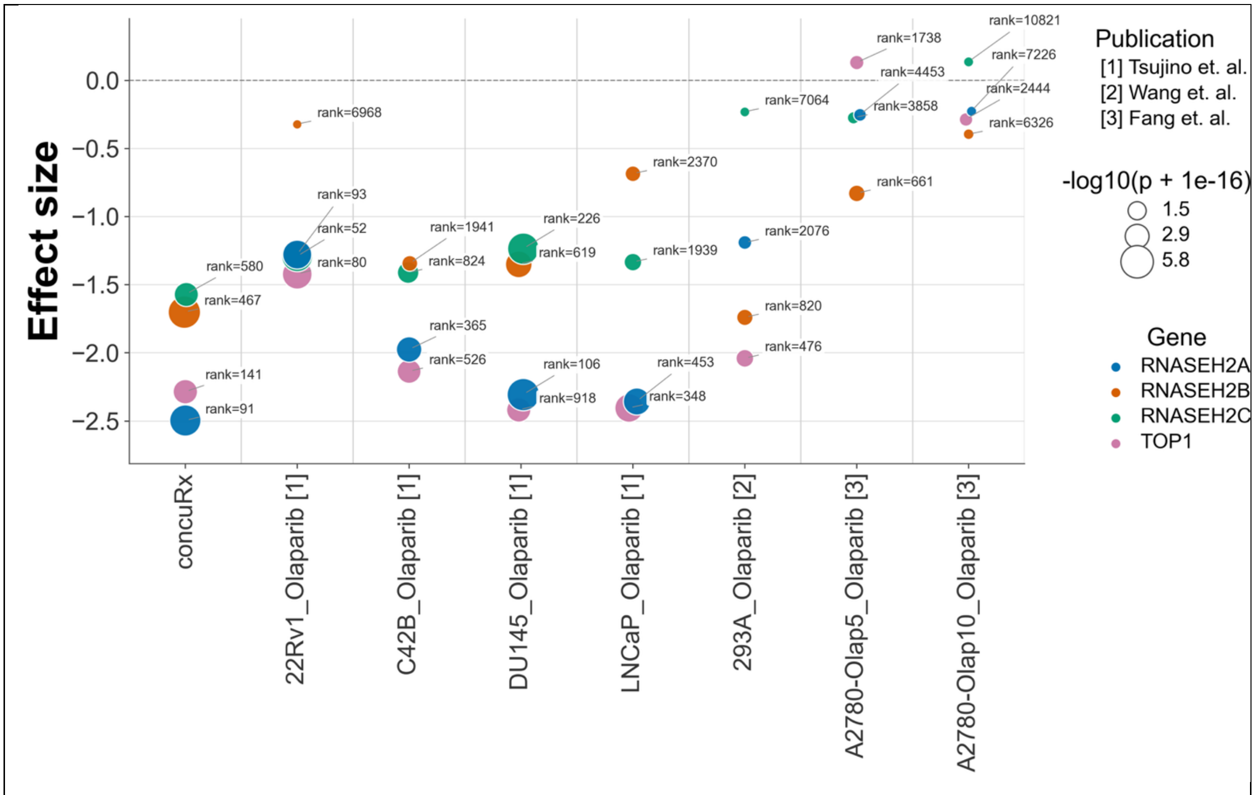


Figure S8: Effect size and significances of RNASEH2 (H12) module genes in PARPi datasets

H13: Fang et al.[4] identified C12orf5/TIGAR as an olaparib-sensitizing gene and showed that TIGAR depletion induces a BRCAness-like state characterized by reduced BRCA1 expression and suppression of Fanconi anemia and homologous-recombination programs, including BRCA2 and RAD51. TIGAR depletion also reduced expression of DNA-replication and cell-cycle genes, including CCNE2, BLM, CDC45, MCM10, MKI67, and PCNA. Importantly, these latter genes were identified primarily through transcriptional changes following TIGAR depletion rather than as top hits from the Fang CRISPR screen itself[4] . TIGAR itself could not be evaluated by ConcuRx because it lacked sufficient representation across the integrated screens. However, ConcuRx independently supported several genetic dependencies associated with the DNA-repair and replication programs affected by TIGAR loss. RAD51 showed sensitizing effects across all seven screen-level inputs and was strongly prioritized by ConcuRx, while CDC45 showed an even stronger integrated sensitization signal. BRCA1, BRCA2, and MCM10 also remained in the sensitizing direction, although with varying statistical support. In contrast, CCNE2, BLM, MKI67, and PCNA showed weaker or heterogeneous cross-study evidence; notably, PCNA was strongly sensitizing in the four Tsujino prostate cancer screens but showed opposing effects in other datasets[5] . Thus, while ConcuRx could not directly assess the TIGAR dependency, it recovered several olaparib vulnerabilities within the HR/replication programs perturbed by TIGAR loss while appropriately reducing confidence for genes with inconsistent cross-study effects.

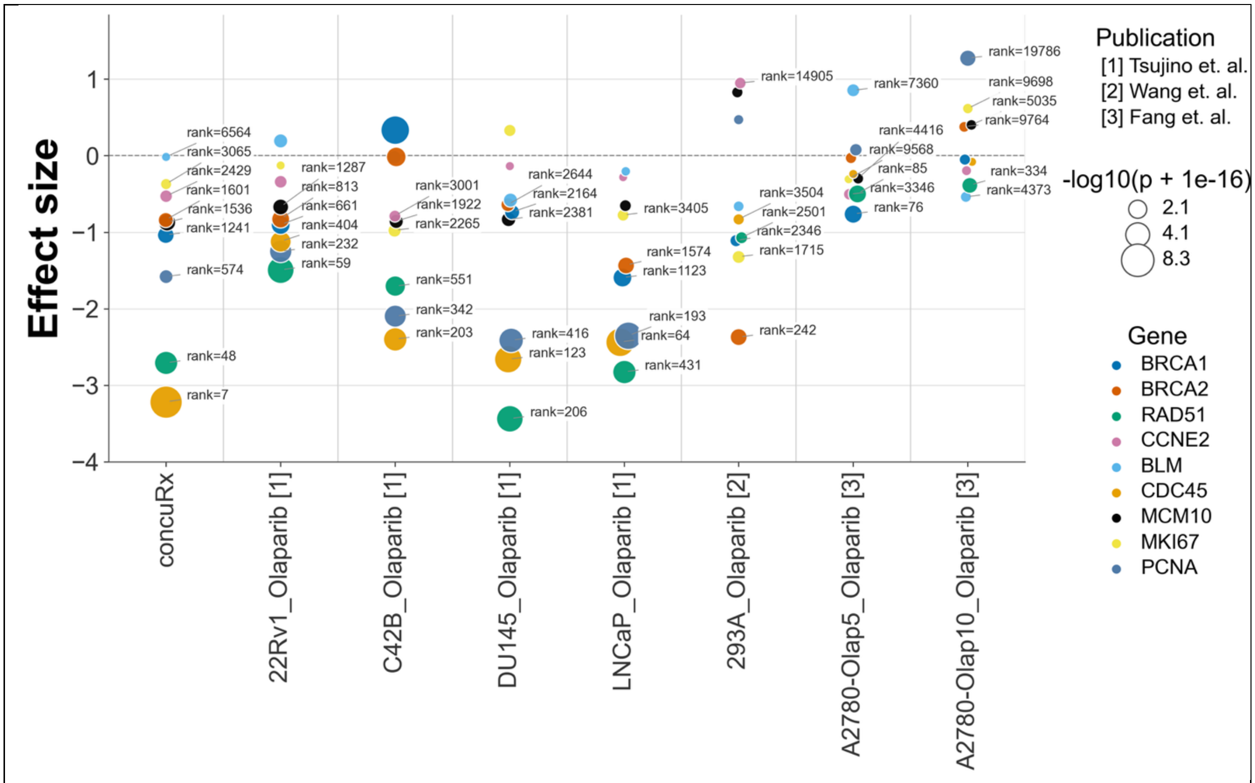


Figure S9: Effect size and significances of H13 module genes in PARPi datasets

H14: Tsujino et al.[5] performed genome-wide CRISPR-Cas9 screens across four BRCA1/2-proficient prostate cancer cell lines to identify genetic determinants of PARPi response. From 67 common negatively selected genes, they selected 10 DDR-associated candidates—MMS22L, ESCO2, RNASEH2B, WDR76, MCM6, POLB, CHD1, FANCI, RAD54L, and HELLS—and validated that deletion of each increased olaparib sensitivity in C4-2B cells. MMS22L was investigated further and shown to sensitize cells to PARP inhibition by impairing RAD51 loading and homologous-recombination repair, with the phenotype dependent on functional TP53. Conversely, loss of CHEK2 promoted PARPi resistance through increased BRCA2 expression and HRR capacity, and combined ATR and PARP inhibition overcame this resistance[5] . ConcuRx recapitulated several of these sensitivity and resistance determinants while revealing substantial cross-study heterogeneity. MMS22L retained a significant integrated sensitizing effect despite variable evidence across individual cell lines, while RAD51—the mechanistic link between MMS22L loss and impaired HRR—showed a strong sensitizing signal. Among the other validated candidates, MCM6 and RNASEH2B showed particularly consistent cross-study sensitization, whereas ESCO2, POLB, RAD54L, CHD1, FANCI, HELLS, and WDR76 showed weaker or more heterogeneous integrated evidence. In the opposite direction, CHEK2 was strongly recovered as a resistance-associated gene, consistent with the experimentally validated resistance mechanism. Thus, ConcuRx recovered key components of both PARPi sensitivity and resistance while distinguishing broadly reproducible dependencies from effects that were stronger in particular prostate cancer contexts.

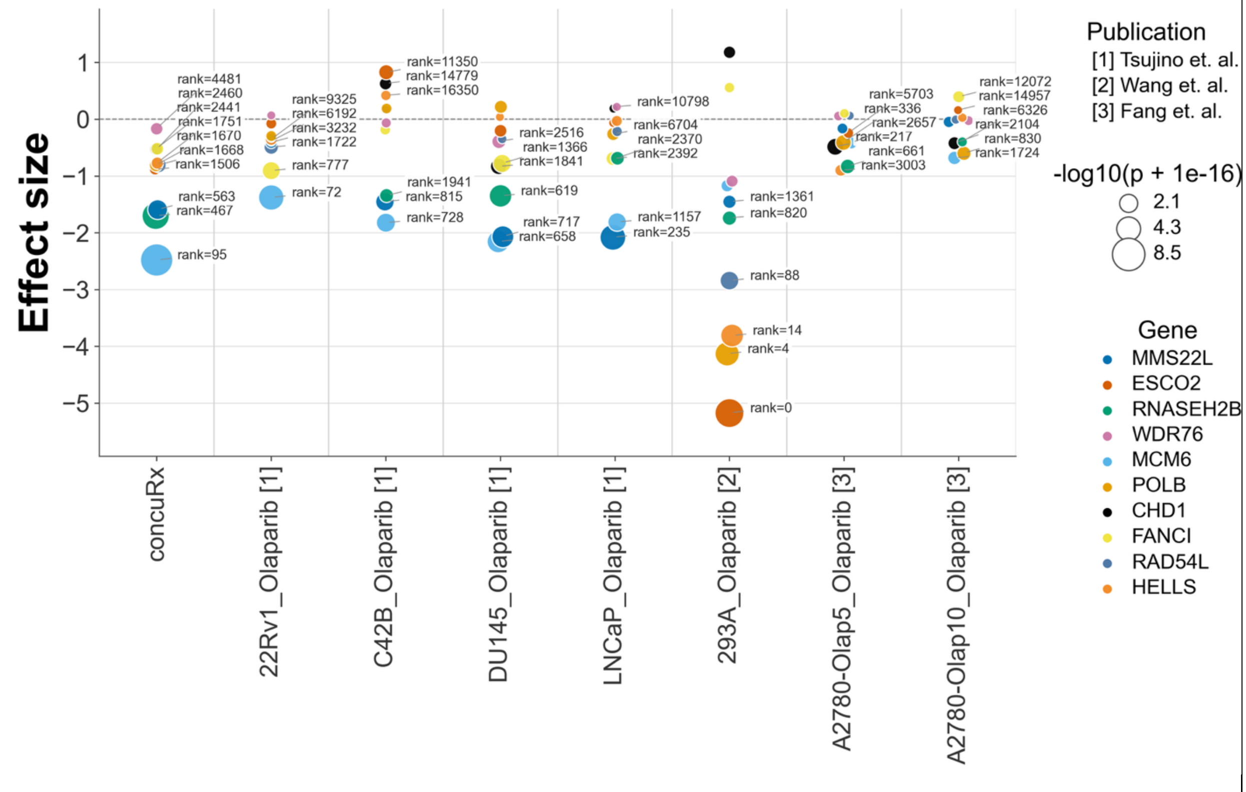


Figure S10: Effect size and significances of H14 module genes in PARPi datasets

H15: Ipsen et al.[6] performed a genome-wide CRISPR-Cas9 screen for genes whose loss confers olaparib resistance in castration-resistant prostate cancer and experimentally validated PARP1, ARH3/ADPRHL2, UBR5, and YWHAE as resistance determinants. Mechanistic studies focused on PARP1 and ARH3, showing that their loss reduces autophagy and that pharmacological induction of autophagy can partially reverse the resistant phenotype. Low ARH3 expression was also associated with prostate cancer recurrence, suggesting potential clinical relevance. ConcuRx independently supported the resistance direction for PARP1 and ARH3, despite the Ipsen dataset not being included in the integration. PARP1 showed one of the strongest resistance effects in ConcuRx, although with substantial cross-study uncertainty, while ARH3 showed a smaller but significant resistance effect. In contrast, ConcuRx did not reproduce the Ipsen resistance direction for UBR5 or YWHAE, which instead showed sensitizing integrated effects, whereas the Ipsen candidates that failed experimental validation—TEAD4, USP48, and CCND1—also lacked ConcuRx evidence of resistance. Thus, ConcuRx independently recovered part of the experimentally validated resistance signal while revealing that some resistance phenotypes may be more context dependent across cell lines and experimental conditions.

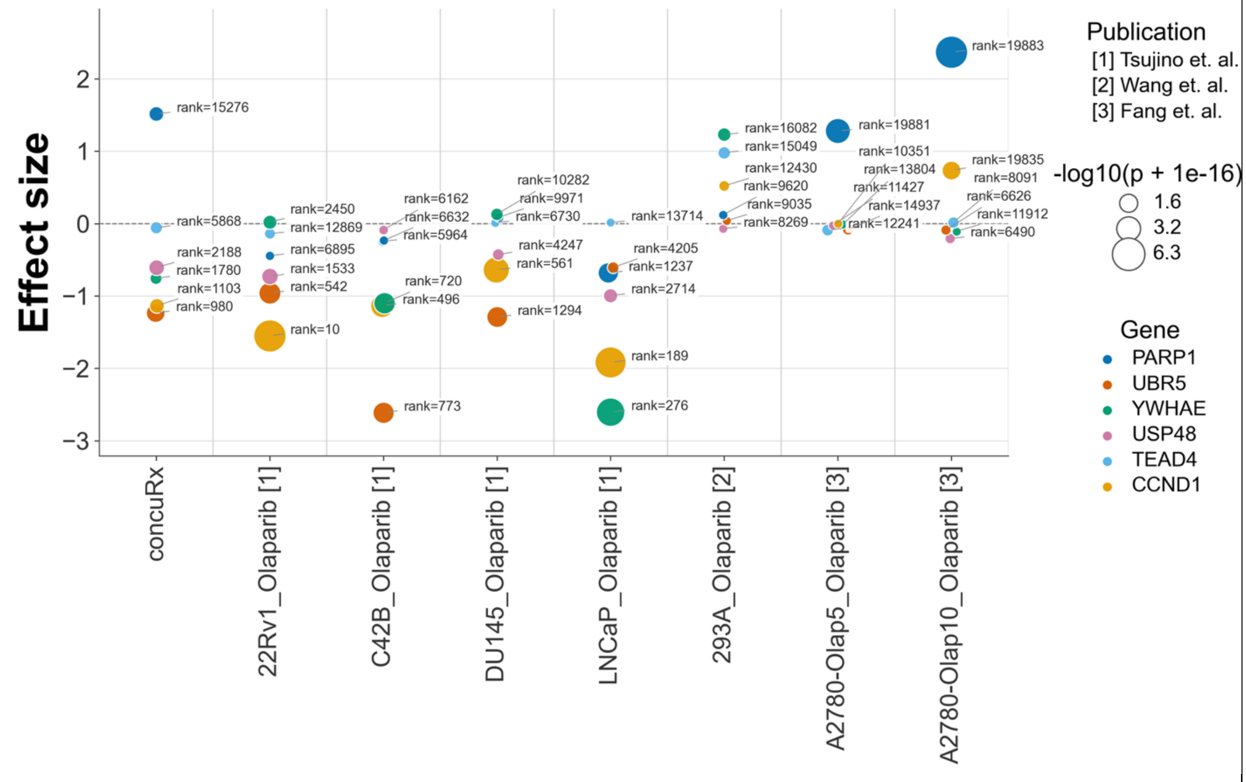


Figure S11: Effect size and significances of H15 module genes in PARPi datasets

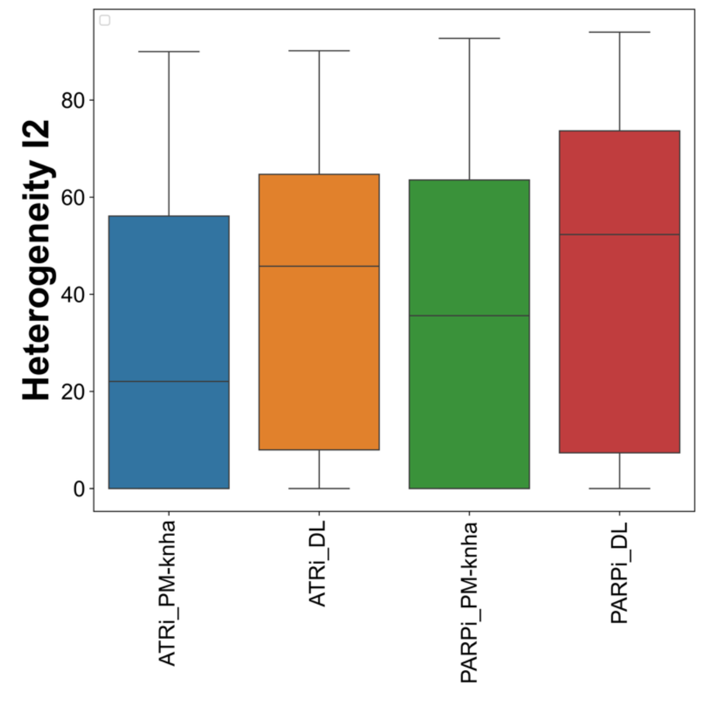

Figure S12: Heterogeneity of the sensitized genes obtained by two random effect meta-analysis (Paule-Mandel or PM-knha and DerSimonian and Laird or DL) in ATRi and PARPi.
